# Fluorescence microscopic evidence for intracellular silicon storage pools in diatoms and their partial separation

**DOI:** 10.64898/2026.11.04.764211

**Authors:** Tobias Reichelt, Julia Breyer, Eike Brunner

## Abstract

Diatoms are unicellular algae that occur ubiquitously in almost every salt- and freshwater habitat on earth. Silicic acid is taken up from the environment and stored in intracellular silicon storage pools (SSPs) to form species-specific silica-based cell walls (frustules). During biosilica formation in the silicon deposition vesicle (SDV), the silicon is transported from SSPs into the SDV.

The present contribution describes the fluorescence microscopic visualization and quantification of silica solely formed from SSPs under silicon starvation. Labeling with the fluorescence marker PDMPO demonstrates continuing cell growth and cell division without silicon uptake from the environment. The SSPs of *Cyclotella nana* may account for up to 1/3 of total cellular silicon and intracellular concentrations up to ca. 540 mM, significantly beyond the saturation level (2 mM).

The chemical state and localization of SSPs within cells are yet unclear. Here, we report a method for subcellular fractionation of *C. nana* by density gradient centrifugation. Quantification of cytoplasmic silicon yields 3–10% of the total cellular silicon corresponding to intracellular concentrations of ca. 50–160 mM. Labeling with NMR-active ^29^Si shows that this silicon occurs as condensed silica, i.e., silicic acid is converted into condensed silica after uptake. However, the detected intracellular silicon concentrations are lower than the value of ca. 540 mM predicted by the fluorescence experiments. It is thus concluded that the missing SSP silicon is either present in the form of large aggregates or compartments. Alternatively, they may also be bound to cell wall biosilica.

## Introduction

Diatoms are unicellular microalgae with a pronounced silicon metabolism. They ubiquitously occur in salt- and freshwater habitats and make up a major part of the phytoplankton. Diatoms are responsible for approximately 40% of the oceanic primary production (Treguer & Pondaven, 2000; P. Tréguer et al., 1995) corresponding to about 20% of the global carbon fixation (Field et al., 1998). Today, about 12,000 diatom species are known (Guiry, 2012). Estimates of the total number range from 20,000 to well over 200,000 species (Guiry, 2012; Malviya et al., 2016; D. G. Mann & Droop, 1996; David G. Mann & Vanormelingen, 2013). For comparison, only about 700 Radiolaria species are known (Biard et al., 2017). This diversity is reflected by a wealth of sometimes bizarre cell wall shapes and patterns. Various geometries and highly ordered structures down to the nanoscale occur. The synthesis mechanisms underlying this wide variety of shapes and the biomolecules involved in biomineralization are still not fully identified and understood. Several polypeptides were isolated from biosilica of diatoms, including silaffins (Kröger et al., 1999, 2001) and silacidins (Wenzl et al., 2008), as well as special long-chain polyamines (Kröger et al., 2000). The latter were found in all diatom species analyzed yet (Sumper et al., 2005) and even in sediments (Annenkov et al., 2018; Bridoux et al., 2011; Bridoux & Ingalls, 2010) as well as the silica-forming *synurales* species *Synura echinulata* (Reinke et al., 2025).

Diatoms actively take up and accumulate silicon (silicic acid) from their environment (Azam et al., 1974; Lewin & Chen, 1968; Lewin, 1955). This silicon is transformed into biosilica during the formation of their cell wall (biomineralization) in a specialized compartment called silicon deposition vesicle (SDV) (Drum & Pankratz, 1964; Heintze et al., 2020; Reimann et al., 1966; Stoermer et al., 1965). Diatoms can store silicic acid in so-called silicon storage pools (SSPs) (Azam et al., 1974; Brunner et al., 2009; Gröger et al., 2008; Kumar et al., 2020; Lewin, 1954; Martin-Jézéquel et al., 2000; Sullivan, 1979; Thamatrakoln & Hildebrand, 2008). However, the intracellular localization and chemical form of SSPs is not yet clear (Martin-Jézéquel et al., 2000).

The decisive causes for the enormous evolutionary success of diatoms are still being debated (Cermeño et al., 2015). In any case, their rapid spread in geological terms, which is still ongoing, seriously impacts global silicon cycles. The silicon concentration in the oceans has changed dramatically over the course of Earth’s history. More than 550 million years ago, the level was close to saturation, i.e., about 10 – 20 times higher than today (Conley et al., 2017; McCutchin et al., 2025). The decline of oceanic silicon concentration coincides with the emergence of the first silicifying organisms (radiolarians, sponges) in the Phanerozoic era (Siever, 1991; P. J. Tréguer et al., 2021). Diatoms appeared much later during the Triassic period about 240 million years ago (Kooistra & Medlin, 1996; Medlin et al., 1997). The earliest fossil record dates back about 190 million years (Sims et al., 2006). During two periods of rapidly increasing biodiversity, first between the Eocene and Oligocene (38 to 32 million years ago) and later in the early Miocene (20 million years ago to the present), the evolutionary and geographical spread of diatoms took its course (Cermeño et al., 2015; Kooistra & Medlin, 1996). Their global success was accompanied by a further decreasing silicic acid concentration in the oceans. A concentration around 600 µM is estimated for time 240 million years ago (Kooistra & Medlin, 1996; Siever, 1991). Today, the oceanic average is approximately 90 µM (P. J. Tréguer et al., 2021). Typical concentrations in surface waters where microalgae grow are about one order of magnitude lower (Cermeño et al., 2015) at an average of <10 µM (Siever, 1991). During the growth period or algal blooms, the environmental silicon concentration can even be close to zero which may explain the need for SSPs.

Lewin investigated the silicic acid uptake of silicon-deficient *Navicula pelliculosa* cultures after silicon replenishment and observed uptake even in the darkness (non-growing conditions). It was hypothesized that silica is deposited on the surface of the pre-existing cell wall (Lewin, 1954). Microscopic observations of the complementary situation showed that cell wall formation (silica deposition on the site of the new cell walls) takes place even in silicon-free culture medium (Coombs et al., 1967). Isotope uptake studies with ^31^Si (Azam et al., 1974; Sullivan, 1979) and ^68^Ge (Azam et al., 1974; Sullivan, 1979) delivered final evidence for the existence of SSPs. It is concluded that the SSPs of some species may be sufficient to build up a complete cell wall and thus account for up to 50% of the total cellular silicon content (tSi_cell_). Examples are *Ditylum brightwellii* and *Conticribra weissffogii* (Chisholm et al., 1978; Martin-Jézéquel et al., 2000). The amount of silicon stored in SSPs depends on the species, environmental conditions, day time, and cell cycle. Estimated intracellular concentrations range from 19 mM (*Cylindotheca fusiformis*), 58–162 mM (*N. pelliculosa*) (Martin-Jézéquel et al., 2000) up to 300 mM (*Lithodesmium undulatum*), 340 mM (*Stephanopyxis turris*) up to even 70–2000 mM (*D. brightwellii*) (Chisholm et al., 1978). These concentrations significantly exceed the saturation concentration of 2 mM for silicic acid at near-neutral pH (Iler, 1979). It is concluded that SSPs must be stabilized to prevent spontaneous and uncontrolled precipitation in the cell (Azam et al., 1974; Sullivan, 1979). Stabilization or complexation of silicic acid by biomolecules and/ or controlled polymerization of silicic acid to oligomers (Annenkov et al., 2008; Azam et al., 1974; S. D. Kinrade et al., 1999; Stephen D. Kinrade et al., 2002; Spinde et al., 2011; Werner & Pirson, 1967), a silica gel (Grachev et al., 2017) or nanoparticles (Brunner et al., 2009; Gröger et al., 2008) have been proposed. Combined transmission electron microscopy (TEM) and energy-dispersive X-ray spectroscopy (EDS) indicated silicon-containing intracellular inclusions in *Synedra acus* (Grachev et al., 2017). Cytoplasmic vesicles of 30–40 nm size fusing with the silicalemma of the SDV in *Thalassiosira eccentrica* were observed by TEM and scanning electron microscopy (SEM). It was hypothesized that these vesicles are silicon transport vesicles (STVs) that originate from the Golgi apparatus (Schmid & Schulz, 1979), although there was no evidence for silicon inside that vesicles. Similar observations were made in *S. turris, D. brightwelli*, and *Gomphonema parvulum* (Dawson, 1973; Li & Volcani, 1985a, 1985b). Consistently high intracellular silicon concentrations were detected in *Cyclotella nana* (formerly *Thalassiosira pseudonana*) by cryogenic focused ion beam (cryo-FIB) in combination with SEM and EDS, leading to the assumption of a homeostatic silicon concentration within the cell. After 12 h silicon starvation, this concentration still amounts to 187 ± 58 mM which even exceeds the value of 140 ± 51 mM determined in cells during exponential growth. The authors report a rather uneven distribution of silicon throughout the cell (Kumar et al., 2020). Solid-state ^29^Si NMR investigations revealed that silicic acid is condensed immediately after uptake and converted into silica. This became evident from ^29^Si pulse labeling of *C. nana* and *D. brightwellii* cultures. Accordingly, the intracellular storage of large quantities of soluble silicic acid (Q^0^/Q^1^) can be ruled out. Instead, it could be concluded that SSPs consist of silica in a pre-condensed form. One hypothesis is that SSPs consist of stabilized silica particles (Bertermann et al., 2003; Brunner et al., 2009; Gröger et al., 2008). Furthermore hypervalent, five or sixfold coordinated silicon species can at least transiently occur in diatoms as shown by ^29^Si NMR studies on *N. pelliculosa* and *C. nana* (Brunner et al., 2009; Stephen D. Kinrade et al., 2002).

The quantitative determination and chemical analysis of SSPs requires their separation from the siliceous cell walls. Established methods are for example the extraction with perchloric acid at 0 °C, trichloroacetic acid at 0 °C, or water at 100 °C (Azam et al., 1974; Sullivan, 1979). These rather harsh methods are, however, likely to change the chemical state of SSPs and to at least partly dissolve cell wall silica. Therefore, mechanical cell lysis by bead beating followed by centrifugation offers a suitable alternative for gentle SSPs extraction (Reichelt et al., 2023). Sensitive silicon quantification can then be performed using the established Molybdenum blue assay (MBA) method (Coradin et al., 2004; Koistinen et al., 2020).

The present work focuses on the development of a density gradient centrifugation-based method with the aim of separating SSPs in *C. nana.* The original and current scientific name is *Cyclotella nana* (Alverson 2011, Poulsen 2023). The former name, *Thalassiosira pseudonana,* is still widely in use. The results of this subcellular fractionation make SSPs accessible for quantification and further chemical analysis. The basic concept is shown in Fig. 1. Moreover, a novel fluorescence microscopic method for SSPs estimation is developed using [2-(4-pyridyl)-5-((4-(2-dimethylaminoethylaminocarbamoyl)methoxy)phenyl)oxazole] (PDMPO) for pulse labeling. Finally, cell cultures are enriched with the NMR-active isotope ^29^Si. Successful separation yields fractions that are enriched with the SSPs and can be analysed by solid-state ^29^Si NMR spectroscopy.

**Figure 1.**
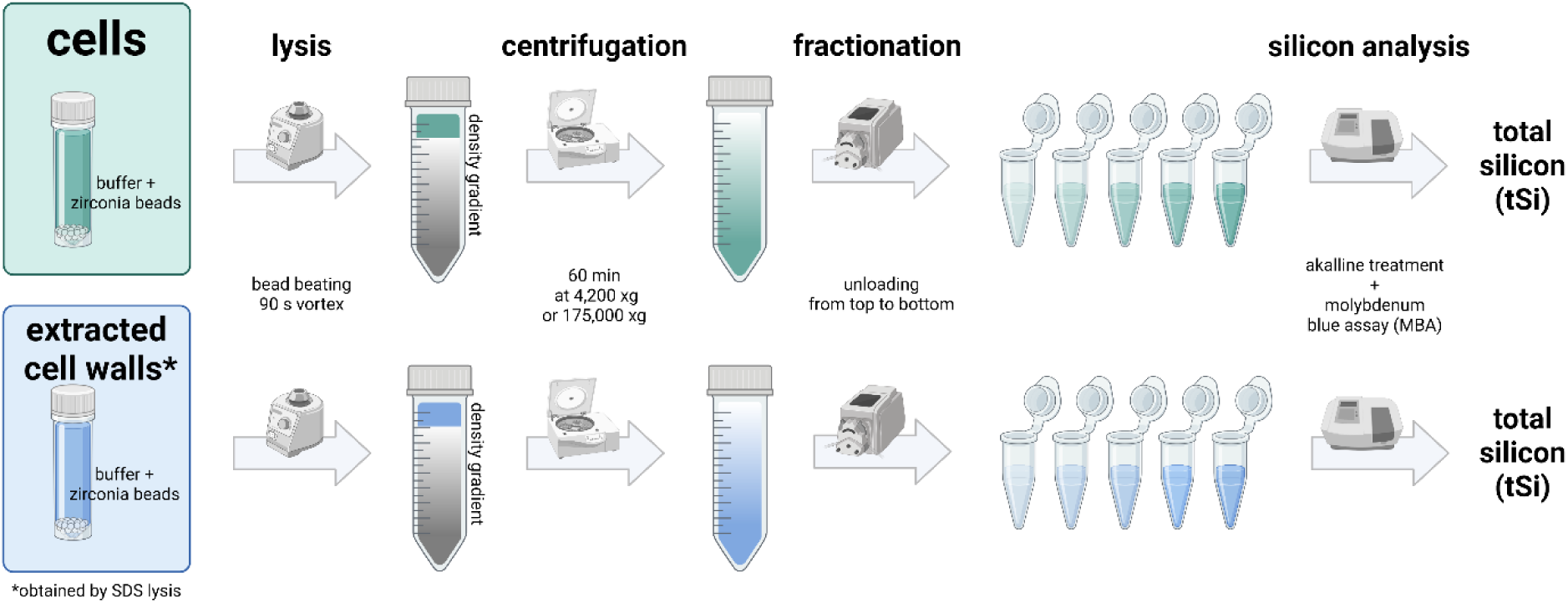
Illustration of the separation process carried out as a parallel experiment to compare cells (top) with extracted cell walls (bottom) as control in order to identify the SSPs in the fractions.

## Materials and methods

### Cultivation of *C. nana* at standard conditions

*C. nana* strain CCMP 1335 was cultivated in sterile, sealed 20-liter polycarbonate containers (Nalgene, Rochester, NY, USA). Artificial seawater (ASW) was prepared with 100 µM of silicic acid (Harrison et al., 1980). For isotope labeling, ^29^Si enriched silicic acid solution was used instead. For silicon starvation and washing, silicon-free ASW (ASW-Si) was prepared accordingly, without the addition of silicic acid. Exclusively plastic labware was used during cultivation. The cultures were maintained at 20 °C in an incubator (RuMed1301, Rubarth Apparate GmbH, Laatzen, Germany) under gentle agitation and constant illumination at 6.000 – 8.000 Lux. The pH was regularly adjusted to 8.0. The cell density was monitored using an automatic cell counter (CellDrop FL, DeNovix, Wilmington, DE, USA) and the silicic acid concentration was measured using the MBA. Harvesting took place during exponential growth and before silicic acid was depleted. This was usually 7-10 days after inoculation at a density not exceeding 0.8·10^6^ ml^−1^. The cells were concentrated on a cellulose filter (round filter, Rotilabo 15A, 2-3 µM retention range, Carl Roth GmbH, Karlsruhe, Germany) and washed with cold ASW-Si under vacuum. To resuspend the cells, the filter was immersed in a small volume ASW-Si, placed on ice and gently shaken for 5 min to gently detach the cells. For silicon starvation the concentrated cell suspension was added to 20 L ASW-Si. For final harvest the suspension was centrifuged (2,500 x g for 5 min at 4 °C), washed with a cold solution of 4.0% sodium chloride and again centrifuged. The pellets were shock frozen and stored at −80 °C.

### Isotope labeling with ^2G^Si

Isotope labeling was carried out using ^29^Si-enriched sodium metasilicate, which was prepared from ^29^SiO_2_ of 98.2% purity (Cortecnet, Les Ulis, France). Equimolar amounts of SiO_2_ and sodium carbonate were mixed in a platinum crucible and heated to 1150 °C in a furnace. For details see the references (Schwarz & Menner, 1924; Wisser et al., 2015). The obtained sodium metasilicate readily dissolves in water and depolymerizes to silicic acid. Insoluble residues were removed by a syringe filter. A yield of 98% was determined with the MBA.

### Quantitative estimation of the SSPs by PDMPO labeling

Methanol extraction was proven by McNair et al. to largely eliminate unbound PDMPO and remove most of the photopigments, which lowers the background fluorescence and increases measurement sensitivity (McNair et al., 2015). Only plastic labware was used during cultivation. Two separate media with 1.5 L of ASW were inoculated with *C. nana* strain CCMP 1335 in sterile and sealed 2.8 L polycarbonate containers (Nalgene, Rochester, NY, USA). A concentration of 0.125 µM PDMPO (AAT Bioquest, Pleasanton, CA, USA) was added to one of these batches for full labeling. Cultivation was carried out under constant illumination (6.000 – 8.000 lux) to exclude cell cycle effects. Harvesting took place during the exponential growth phase after 5 days at a cell density of 0.6 - 0.8 · 10^6^ ml^−1^. The fully labeled cells were harvested by centrifugation (2,500 x g for 5 min at 4°C). The pellet was suspended in 20 ml methanol (HPLC grade) and left for extraction at 4°C for at least 24 h. Methanol extraction was proved by McNair et al. to largely eliminate unbound PDMPO and remove most of the photopigments, which lowers the background fluorescence and increases measurement sensitivity (McNair et al., 2015). The unlabeled cells were harvested with a cellulose filter (round filter, Rotilabo 15A, 2-3 µM retention range, Carl Roth GmbH, Karlsruhe, Germany) and washed to completely remove silicic acid. To achieve this requirement, the cell suspension was filtered until less than 100 ml remained above the filter. Then 500 ml of ASW-Si was added for washing and filtered until less than 100 ml remained. The washing step was repeated 6 times. After these steps, the remaining Si concentration is negligible, i.e., it is equal to the trace contamination of silicon in ASW-Si (results section). The unlabeled cells were transferred directly from the filter into fresh ASW-Si (silicon starvation) containing 0.125 µM PDMPO (pulse labeling). After further incubation for 48 h at continuous illumination, the cells were harvested by centrifugation as described before and suspended in methanol.

The extracted cells were washed once with methanol and twice with 4% (w/v) sodium chloride solution (2,500 x g for 5 min) before the analysis under the microscope. Aliquots of the sample were pipetted onto a microscope slide and covered with a cover slip. Adding sucrose to the suspensions to increase the viscosity prevented the cells from moving and mechanical damage. Images were taken in the fluorescence channel of the microscope Biozero BZ8000 (Keyence, Osaka, Japan) at a wavelength of 360 nm for excitation and 535 nm for emission at 20fold magnification and an acquisition time of 10 s. No oversaturation of the detector occurred for these parameters. At least 6 observations were made in z-stack mode. A stack of 6 to 12 individual images with a vertical focus distance of 1-2 µm was sufficient to focus on all cells in the layer. The images were calculated to form a resolved image (full focus) using the analysis software of the manufacturer (Keyence). The intensities were quantified using the image analysis software *ImageJ*. Different from the original method introduced by Mc Nair et al., no fluorescence standard was used in the present study. Instead, the PDMPO fluorescence intensity of the pulse labeled cells was directly correlated to the intensity of the fully labeled cells. Additional species were investigated: *C. weissffogii* strain CCAP 1085/18, *N. pelliculosa* strain SAG 1050-3*, Cyclotella cryptica* strain CCMP 332 *and Biddulphia tridens* unknown strain (see supporting information, Figs. S2 – S5 and Tab. S1).

### Bead beating lysis

The samples, dense cell pellets or biosilica, not exceeding 300 µL in volume, were given in 2.0 ml microtubes. The cold lysis buffer was added to a final volume of 1500 µL. This buffer at pH of 7.5 contained 150 mM sodium chloride, 50 mM 4-(2-hydroxyethyl)-1-piperazineethanesulfonic acid (HEPES), 1 mM mercaptoethanol (ME), and 100 µM phenylmethyl-sulfonyl fluoride (PMSF). ME and PMSF were added shortly before the lysis. Following the addition of 2.2 g yttria-stabilized zirconia (YSZ) beads of 0.1-0.2 mm diameter (Sigmund Lindner GmbH, Warmensteinach, Germany), the microtube was vigorously mixed on a vortex mixer for 90 s with breaks for cooling. The lysate was separated from the beads and kept at 4 °C. (Reichelt et al., 2023).

### Density gradient centrifugation

A continuous sucrose gradient was prepared from 10% (w/w) and 60% (w/w) sucrose solutions using a gradient mixer, a peristaltic pump, and a 14G cannula attached to the bottom of a 50 ml centrifuge tube. The solutions were buffered with 20 mM HEPES at pH 7.5. The final gradients were cooled until the lysate was carefully layered on top using a syringe and a cannula. Centrifugation at 4,200 x g (5,000 rpm) and 4°C was performed for 60 min with acceleration and deceleration set to low values. The tube was emptied beginning from the top with a canula and a peristaltic pump and the collected fractions were immediately frozen. The linearity of the density gradient was confirmed by measuring the refractive index of the fractions.

### Isopycnic density gradient ultracentrifugation

Ultracentrifugation was performed with unlabeled samples and isotope labeled samples. A pellet of cells or extracted cell walls of 2000 µL volume (ca. 13 x 10^9^ cells) was used for the experiment. The lysis resulted in 10 ml of lysate, which was layered over a stepped density gradient covering a density range of 1.11 - 1.43 g/ml. The seven layers, each of 4 ml volume, consisted of an aqueous solution of iodixanol (OptiPrep, 60% w/v aqueous solution, Serumwerk, Bernburg, Germany) in the upper part and iohexol (Nycodenz, powder, Serumwerk, Bernburg, Germany) in the lower part of the column, or a mixture of the two components. The upper five layers contained 20, 30, 40, 50 and 60% (w/v) OptiPrep and sodium chloride at concentrations of 122, 108, 93, 79, 72 and 65 mM to obtain iso-osmolar conditions. The sixth layer consisted of a 1:1 mixture of OptiPrep and Nycodenz at 70% (w/v) at ca. 440 mOsm, and the seventh layer contained 80% (w/v) Nycodenz at ca. 770 mOsm. All solutions were buffered with 20 mM HEPES at pH 7.5.

Ultracentrifugation was performed in open-top thin-wall ultra-clear polycarbonate tubes (Beckmann Coulter, Brea, CA, USA) with a volume of 38.5 ml (25×89 mm) using the rotor SW 32Ti (Beckmann Coulter) on the ultracentrifuge Optima XE90 (Beckmann Coulter) at 32,000 rpm and 4 °C for 60 min. The relative centrifugal force (RCF) was thus 175,000 x g at the bottom and 76,600 x g at the top of the tube. Acceleration and deceleration were set to intermediate values. The tube was emptied beginning from the top using a cannula and a peristaltic pump. The fractions were removed in a way that the transition between two density layers was exactly in the middle of the fraction and the particles in the boundary area were completely taken up. Only the first fraction (8 ml) and the last fraction with the pellet (2 ml) not contained a density transition. The fractions were carefully homogenized and samples were taken for the analysis of the total silicon amount (tSi). The fractions were shock-frozen and stored at – 80 °C. Selected fractions were analyzed by NMR spectroscopy.

### Alkaline treatment

For the alkaline treatment, the silicon concentration in the sample (solution or suspension) should not significantly exceed 2 mM. Otherwise, it should be diluted before. 0.2 M sodium hydroxide solution was added to the sample with a volume at least equal to the volume of the sample. The mixture was heated to 95 °C for 15 min. After cooling, the same volume of 0.2 M hydrochloric acid was added to the mixture for neutralisation. The tube is filled with water to the desired volume and dilution (Koistinen et al., 2020).

### Molybdenum blue assay

An aliquot of the alkaline treated sample was transferred to a 2.0 ml microtube. For the standard addition, a series of 5 aliquots was taken. 4 aliquots were mixed with a silicic acid standard solution (BerndKraft, AnalytiChem GmbH, Duisburg, Germany) to a final added standard concentrations of 5, 10, 15 and 20 µM (based on the final volume of 1800 µL). The microtubes were filled with ultrapure water to a volume of 1620 µl. 60 µL of a solution of 81 mM ammonium heptamolybdate and 1.8 M sulfuric acid was added. After mixing and equilibration for at least 3 min, 60 µL of 1 M oxalic acid solution was added, followed by mixing and 1 min equilibration. Finally, 60 µL of 100 mM ascorbic acid solution was added. Measurements were performed after 30 min. If precipitation occurred or particles were present, the microtube was centrifuged at 20,000 x g for 3 min. The supernatant was transferred to a cuvette (10 mm, polystyrene, VWR, Radnor, PA, USA) for photometric measurement (Cary 60, Agilent, Santa Clara, CA, USA). The background of a blank sample was subtracted. (Koistinen et al., 2020). The silicic acid concentration in the cultivation media was quantified accordingly. The trace contamination of ASW-Si was determined using diluted silicic acid standards with final added concentrations of 0.2, 0.4, 0.6 and 0.8 µM. Multiple blanks in separate cuvettes (n=16) were measured to determine the limit of quantification (LOQ) of 0.17 µM. The absorbance was detected for 5 s (average time) at a wavelength of 810 nm.

### Solid-state ^29^Si NMR spectroscopy

The ^29^Si-enriched samples were investigated without drying. The fractions (4 ml) obtained after ultracentrifugation were suspended in cold lysis buffer (8 ml) to reduce the solution density, followed by centrifugation at 4,200 x g and 4 °C for 40 min. Centrifugation was repeated with the supernatant for 90 min, to recover few amounts of additional material. The wet pellets of the washed fractions and the wet pellets of the whole cells were resuspended with a few drops of the buffer. The fractions belonging to the gradient densities of 1.21 g/ml and 1.26 g/ml were completely combined to obtain sufficient sample material for the measurement. The suspension particles were transferred to a rotor insert (disposable insert for 4 mm magic angle spinning (MAS) rotor, 25 µL nominal volume, Bruker BioSpin, Karlsruhe, Germany) by repeated centrifugation. To this end, a 200 µl pipette tip was cut on the narrow end to tightly fit inside the orifice of the rotor insert. Both were connected with a rubber tubing cut short. The wide end of the tip was cut so that the whole construction fit in a 2 ml Eppendorf tube. The insert and the tip were filled with up to 80 µL of the suspension and were centrifuged for 10 min starting at 1,000 x g and slowly rising the RCF to 20,000 x g. The supernatant was carefully removed and additional suspension was added. The step was repeated until the particles of the suspension were completely transferred. In case the suspension spilled out it helped to reduce the volume in the tip. The sample was tightly compressed with the screw cap of the insert and given into a 4 mm MAS rotor for solid-state NMR spectroscopy. Spectra were recorded by direct ^29^Si excitation at a sample spinning rate of 10 kHz within 1–4 days according to 250–1000 scans with a recycle delay of 360 seconds on a BRUKER Avance 300 MHz spectrometer using a 4 mm double resonance {H,X} MAS NMR probe (Bruker BioSpin, Karlsruhe, Germany).

### Determination of the average radius, volume and silicon amount of *C. nana*

Cultures of *C. nana* in the exponential growth phase were concentrated by centrifugation and washed several times with ASW-Si to remove dissolved silicic acid. Aggregation must be avoided during centrifugation and resuspension, since aggregates distort cell counting. The cell density (n > 16) was determined on the automated cell counter. The suspension was alkali-treated and tSi was determined using the MBA and the standard addition method (n = 7). The determined average silicon content of a cell in exponential growth was 138 ± 5 fmol (95% confidence interval (CI), n = 3). For volume determination, the cell size was determined from microscopic images using *ImageJ* software. The average radius amounted to 2440 ± 46 nm (95% CI, microscopically determined as the average over n = 414 cells). The mean volume of a cell in exponential growth was 88.1 ± 3.9 fL (95% CI, microscopically determined as the average over n = 204 cells) assuming a cylindrical cell shape.

## Results

### Quantification of SSPs by PDMPO pulse labeling

In order to visualize and estimate the silicon amount in SSPs of *C. nana*, the following indirect approach was applied (cf. Fig. 2, top). Cells in the exponential growth phase are transferred into silicon-free artificial seawater (ASW-Si) and pulse labeled with PDMPO. After 48 h of incubation, fluorescence microscopy reveals new biosilica formed after pulse labeling whereas previously formed biosilica is not labeled because PDMPO is only incorporated during biogenesis/ silica precipitation (Shimizu et al., 2001). Since, however, no external silicon is available, biosilica synthesis can only occur at the expense of SSPs present before pulse labelling and the fluorescence intensity thus reflects this SSP-derived biosilica.

**Figure 2.**
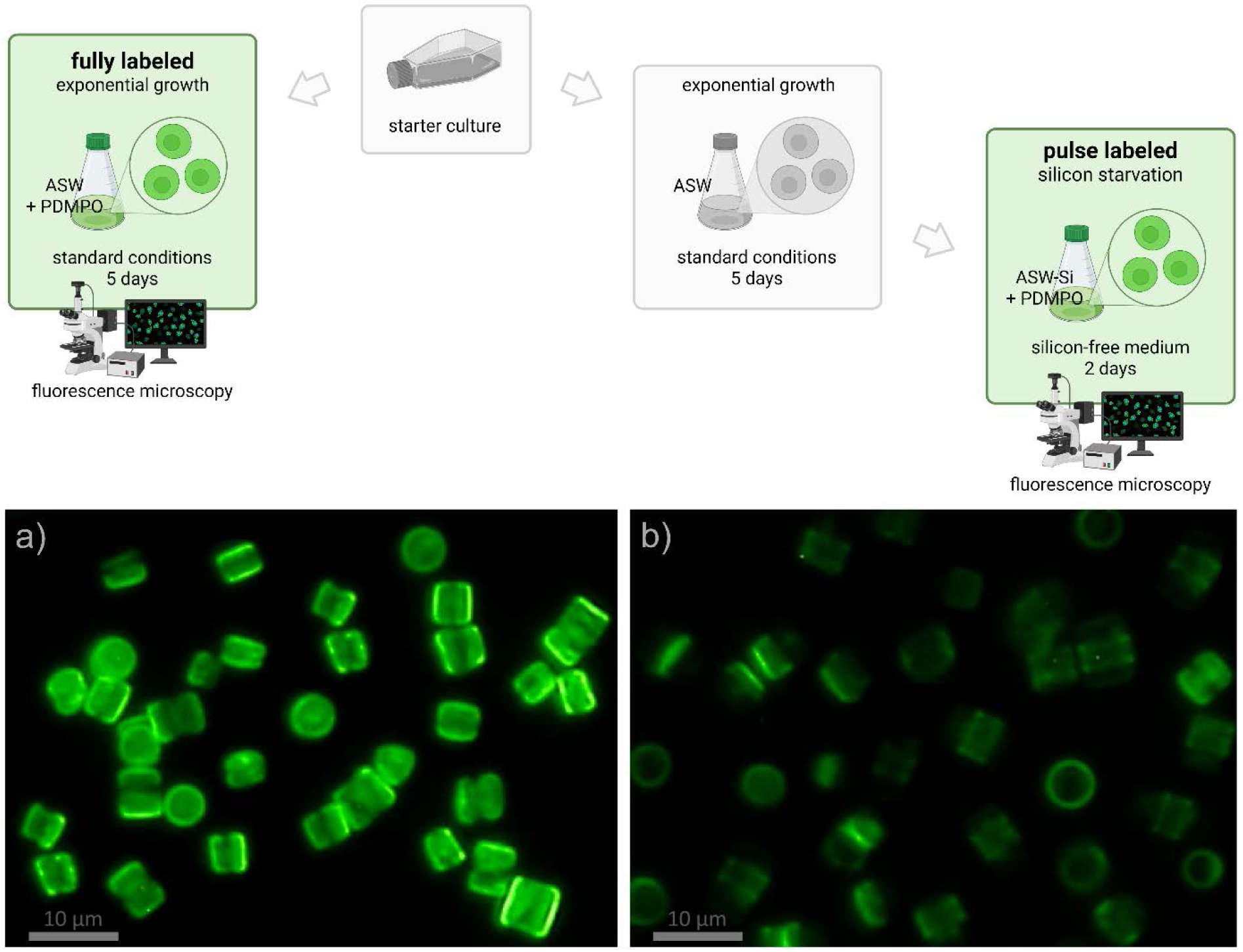
Top: Scheme illustrating the experiment for quantifying the SSPs based on fluorescence intensities of PDMPO in cellular biosilica after full labeling (left) and pulse labeling in silicon-free medium (right, ASW-Si: Si-free ASW). Bottom: Microscopic images show the fluorescence of PDMPO in the biosilica of methanol extracted cells of *C. nana* after full labeling (a) and pulse labeling in silicon-free medium (b).

Remarkably, PDMPO pulse labeling results in detectable fluorescence for more than 99% of the cells (Fig. S1) after 48 h in silicon-free medium (Fig. 2b). Cell division is recognizable by the presence of labeled valve pairs. For quantification, a cell culture in the exponential growth phase was fully PDMPO labeled in Si-containing artificial seawater (ASW) and used as reference (Fig. 2a).

The cells were extracted with methanol prior to microscopic imaging to remove PDMPO that is not incorporated into biosilica (McNair et al., 2015). The fluorescence intensities were determined using the software *ImageJ*. Cell counting was performed manually. To estimate the SSPs, the quotient of the mean fluorescence intensities of the pulse labeled cells (Si starved) and the fully labeled cells (exponential growth) was calculated. The obtained result of 34 ± 6% thus represents the average fraction of SSP-silicon relative to the total silicon amount of a cell (tSi_cell_) in exponential growth. Six images with an average of 90 cells were evaluated from each sample. Note that the experimental error (95% CI) represents the average fluorescence intensity variation of a cell in these images (n = 6), it does not reflect the variability within the cell culture. Also note that the applied fluorescence technique would only detect the entire amount of SSPs after their complete utilization by the cells. To come close to this situation, we have chosen 48 h growth period for the experiments and observed fast growth only within 24 h, at maximum 48 h after transfer to ASW-Si. Afterwards, cell growth slows down severely or stops completely.

The described estimation of SSPs in *C. nana* relies on the observation that PDMPO incorporation into cell wall biosilica, also referred to as biogenic silica (bSiO_2_), occurs with constant molar bSiO_2_ : PDMPO ratio - independent of the external silicic acid concentration in the medium. The fluorescence intensity can thus be used as a quantitative biosilica marker (Leblanc & Hutchins, 2005; McNair et al., 2015). However, there is also preliminary work indicating an increasing molar bSiO2 : PDMPO ratio if the external silicon concentration falls below 3 µM (McNair et al., 2015). This would lead to an overestimation of SSPs and the determined value of 34% should then be considered as an upper bound. However, the observation of an increasing bSiO_2_ : PDMPO ratio for very low Si concentrations (< 3 µM) suffers from an inherent methodical problem of the ^32^Si-labeling experiment as discussed below in detail (discussion section).

In our experiments, a fluorescence standard - the fully PDMPO-labeled cells - were used as reference. Therefore, we obtain the dimensionless fraction representing the SSPs in relation to the total cellular Si amount tSi_cell_ (see above). In addition, this experimental setup ensures that species-specific variability of the molar bSiO2 : PDMPO incorporation ratio has no effect. From this fraction, the Si concentration of SSPs [tSi_SSP_] = 540 ± 100 mM and the corresponding cellular Si amount of tSi_SSP_ = 47 ± 8 fmol/cell were determined using the measured total cellular Si amount tSi_cell_ = 138 ± 5 fmol and a cell volume of 88.1 ± 3.9 fL (Tab. 1) (materials and methods). Four additional diatom species have been analyzed with this fluorescence method, their estimated SSPs range from ca. 4% to 33% of tSi_cell_ (Tab. S1 and Fig. S2 – S5).

**Table 1.**
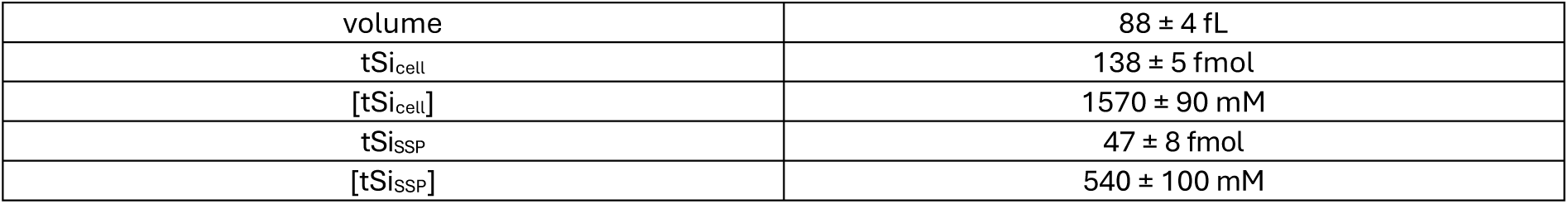
*C. nana* cell parameters: Average cell volume, total cellular silicon amount tSi_cell_ and concentration [tSi_cell_] determined using the Molybdenum blue assay. The silicon amount tSi_SSP_ and concentration [tSi_SSP_] of SSPs was estimated based on the described fluorescence method. The experimental error is given as 95% CI.

ASW-Si was tested for silicon contaminations using the MBA and standard addition. A trace concentration of 0.26 ± 0.2 µM Si was found. The LOQ was determined to be 0.17 µM (materials and methods). In order to minimize the influence of that contamination, the cell suspensions were adjusted to a high density before silicon starvation (ca. 1.0 · 10^6^ cells/ml for C. nana). As a result, the contamination would account for ca. 0.26 fmol silicon per cell. Compared to the tSi_cell_ of ca. 138 fmol (Tab. 1), the influence of the contamination is negligible.

In this context, it should be mentioned that the physical dissolution of pre-existing biosilica is considered as a potential undesired source for silicon release. As far as we know, there is no data available on how fast this process occurs on living cells. We considered dead cells to be a potential pathway for silicon release. A low dead cell count of < 1% was thus ensured before the starvation experiment.

It should be emphasized, that the obtained value of 34% is considered as an upper bound of the SSPs proportion on tSi_cell_. An important issue is the uncertainty about the precise ratio of bSiO_2_ : PDMPO in the biosilica synthesized during silicon starvation, as discussed above and in detail in the discussion section. Furthermore, it is not certain that SSPs are indeed stainable with PDMPO in the exponentially growing cells (control), since the chemical state of SSPs is still not fully clear. In that case, the resulting SSPs would be reduced to 25% of tSi_cell_.

### Density gradient centrifugation

A continuous sucrose density gradient was used to fractionate exponentially grown cells after bead beating cell lysis. For control, extracted cell walls were treated and analysed in parallel. Centrifugation was performed for 60 min at 4,200 x g and 4°C. With that parameters separation is predominantly based on particle size. The total amount of silicon tSi in the obtained fractions was determined after alkaline treatment with MBA using the standard addition method (n = 5) (materials and methods). Fig. 3 shows the tSi-values of the individual fractions in percent. These proportions refer to tSi of the entire sample (tSi_sample_), i.e., the sum of all fractions. Since tSi_sample_ is just a multiple of the average silicon amount of a cell tSi_cell_, the proportion of the fractions on tSi_sample_ is equal to their proportion on tSi_cell_.

**Figure 3.**
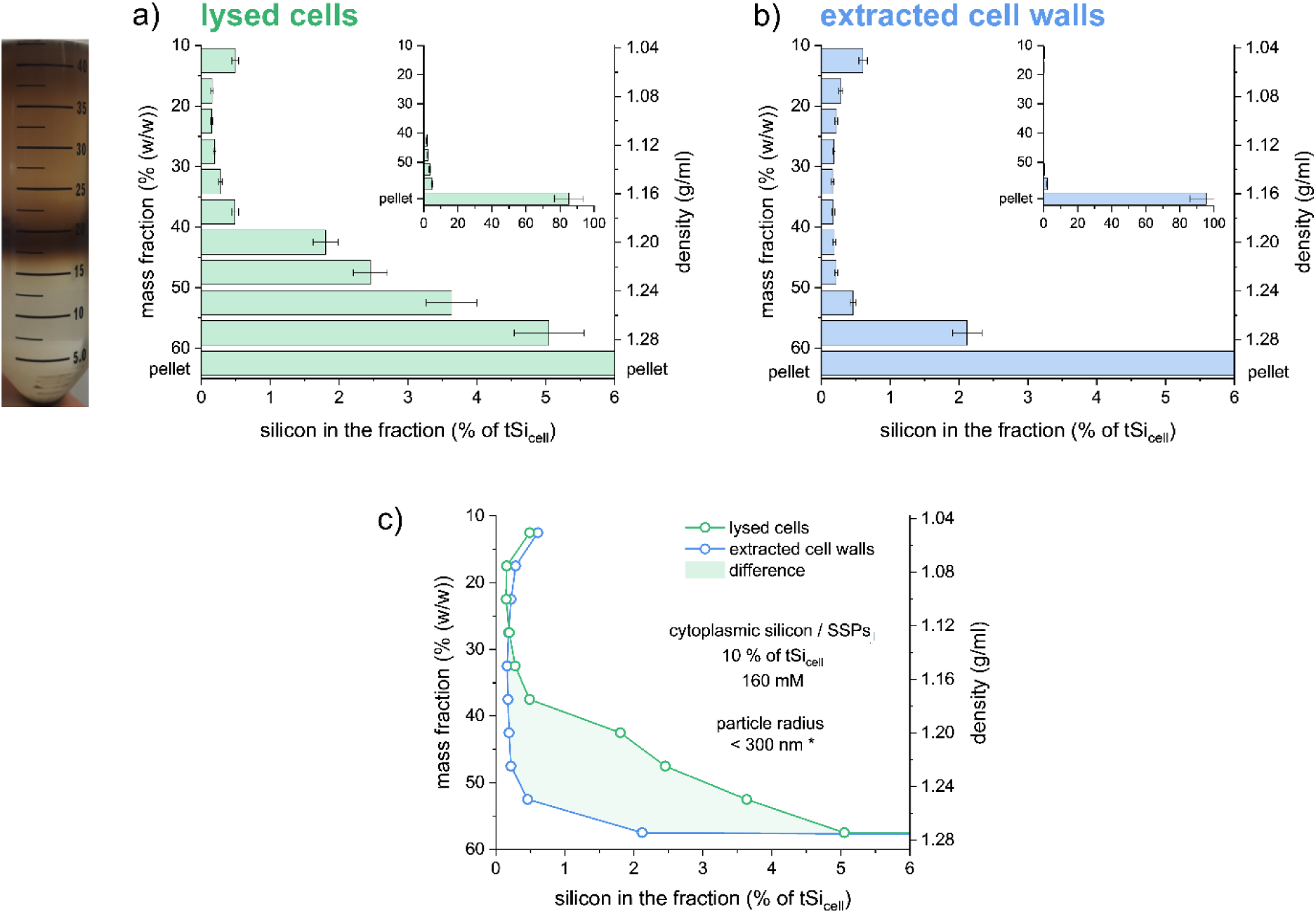
Top, left: Photograph of the sucrose density gradient with cell lysate after centrifugation. Diagrams showing the tSi-values of the different fractions for the cells (a) and the extracted cell walls (b) as fractions of tSi_cell_. c) Difference between the curves for cells and extracted cell walls. This difference represents an estimate for the cytoplasmic silicon, which is assumed to be part of the SSPs. * Upper bound radius for pure silica particles (2.20 g/ml) calculated with the Stokes’ law. Larger silica particles – which may be part of the SSPs as well – will be found in the pellet (Fig. S6 and App. S1). Error bars: 95% CI.

Extracted cell walls after bead beating treatment (Fig. 3b) behave pronouncedly different from the cell lysate. They almost completely sediment in the pellet (95.4%) and only 2.1% are found in in the fraction directly above the pellet. Lysed cells (Fig. 3a) exhibit significantly higher silicon contents especially in the fractions beyond a density of ca. 1.18 g/ml. This increase starts below the dark visible band containing the chloroplasts (Fig. 3, top left). The reference sample (extracted cell walls) indicates that only limited amounts of cell wall fragments are found in the fractions on top of the pellet. This approach of using extracted cell walls as control has already been applied in our previous study (Reichelt et al., 2023). In the present study, it indicates once again that silicon-containing particles (any combination of silicon with organic material of unknown constitution and chemical form) are present in the cytoplasm. By calculating the difference between the quantities found in the gradient of the cell lysate on the one side and the extracted cell walls (control) on the other side, it is concluded that these silicon-containing particles make up 10 ± 0.7% of tSi_cell_ in the lysed cells. The conclusion that those silicon-containing particles are likely originating from the cytoplasm and not the cell walls, follows from the fact that cell wall fragments (control) are sedimented almost completely and are missing in the gradient above the pellet, whereas the cell lysate shows high silicon proportions in the gradient.

### Isopycnic density gradient ultracentrifugation

To further improve the separation of SSPs, ultracentrifugation was performed with media of high density. In order to obtain (near)isopycnic condition, high RCF was applied. In this case separation is based solely on the particle density. A stepped gradient was used consisting of isoosmolar OptiPrep (iodixanol) solutions up to 1.32 g/ml density. Nycodenz (iohexol) was then used for densities of 1.37 and 1.43 g/ml at hyperosmolar conditions. Centrifugation was performed at 175,000 x g and 4°C for 1 h (materials and methods). According to calculations, isopycnic conditions apply to cytoplasmic particles with a radius exceeding ca. 30-50 nm (Fig. S7). Such particles reach equilibrium (buoyant density) after 60 min centrifugation. Silica has a density well above the gradient (ρ_biosilica_ ≈ 2.2 g/ml). Consequently, there is no equilibrium state in the gradient for pure silica particles. Calculations show that silica particles reach the pellet within the runtime of 60 min, if their radius is greater than 40 nm. Below that threshold value, the particles remain in the gradient (Fig. S7). The tSi-values of the various fractions were determined by MBA after alkaline treatment using standard addition (n = 5) for calibration (materials and methods). After ultracentrifugation of lysed cells (Fig. 4a), 92.0% of the silicon was found in the pellet and 8.0 ± 0.4% in the fractions of the gradient. A significant amount of tSi representing 5.8 ± 0.4% of tSi_cell_ was found in the upper half of the gradient in the density range of 1.01–1.27 g/ml (Fig. 4a), whereas the control of extracted cell walls only contained 0.48 ± 0.05% in this range (Fig. 4b). Consequently, almost no cell wall fragments are located in this density range and the observed difference of 5.3 ± 0.4% (relative to tSi_cell_) thus represents silicon species being part of the SSPs (Fig. 4c). The silicon found at densities higher than 1.27 g/ml, in the density range of 1.32–1.43 g/ml, mainly consists of cell wall fragments, as shown by the control. The concentrations are significantly lower than in the fractions above. The presumed SSPs from the cytoplasm are therefore clearly separated from the cell walls, that are predominantly found in the pellet.

**Figure 4.**
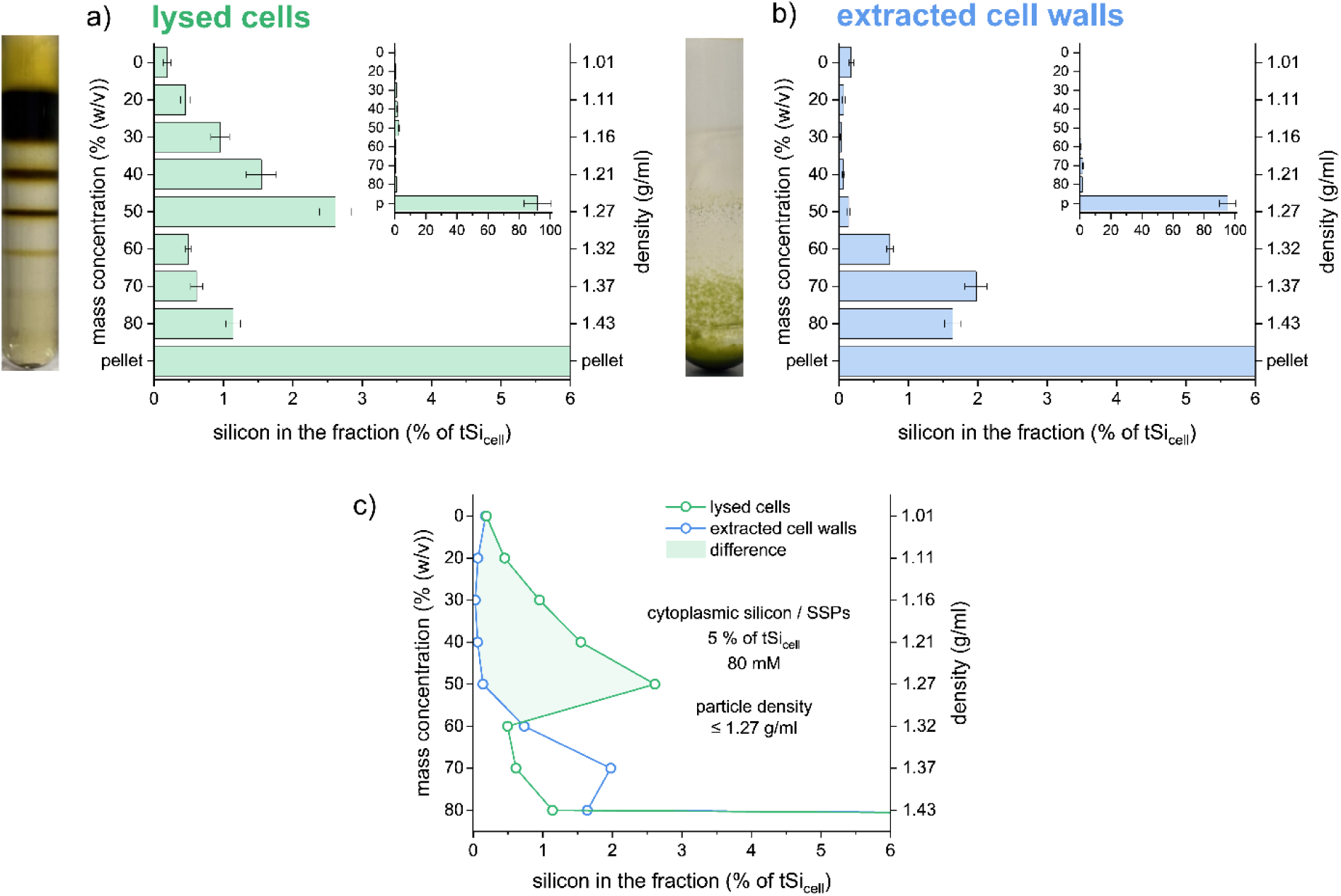
Top: Photographs of the density gradients after ultracentrifugation and diagrams showing the corresponding tSi in the lysed cell fractions from cultures in exponential growth (a) and the control experiment with extracted cell walls (b). tSi-values are fractions of tSi_cell_. c) Difference between the curves for lysed cells and extracted cell walls providing an estimate for cytoplasmic silicon which is assumed to be part of the SSPs (Fig. S7 and App. S1). Error bars: 95% CI.

The highest contents of the cytoplasmic silicon are found at densities of 1.21–1.27 g/ml. It is concluded based on calculations that the silicon-containing SSP constituents located in that fractions are in equilibrium state and have a density very close or equal to the density of the gradient in this region. This follows from the fact that particles of a higher density, such as pure silica, would need to have very small radii of approximately 10–16 nm to reach this region of the gradient (Fig. S7). However, such particles would have been detected in the sucrose gradient of the previous investigation, namely in the uppermost fraction with the lowest density (Fig. S6). As already shown, there is no cytoplasmic silicon found in this fraction (Fig. 3c) and consequently such particles do not occur in a pure and free form.

### ^29^Si NMR spectroscopy

For ^29^Si solid-state NMR analysis, ^29^Si-enrichment was performed by cultivation of *C. nana* in a growth medium with ^29^Si-enriched silicic acid under standard conditions. After the harvest of the exponentially grown cells, half of the culture was transferred to a silicon-free medium and incubated for 24 h to obtain silicon starved cells (materials and methods).

The solid-state ^29^Si MAS NMR spectrum of the exponentially grown whole cells (Fig. 5b) shows the characteristic signals of biosilica representing Q^n^ groups (Fig. 5c). The exponent n indicates the degree of polycondensation: n = 0 for monosilicic acid Si(OH)_4_ and n = 4 for completely polycondensed silica SiO_2_. In the spectrum, the Q^4^ signal occurs with the highest intensity, followed by Q^3^. The group Q² is hardly detectable. This is in line with the signals typically observed in biosilica (Belton et al., 2012; S. Mann et al., 1983). Q^1^ and Q^0^ do not appear in the solid-state NMR spectrum (Fig. 5b). However, Q^1^ and Q^0^ are the predominant signals in the liquid-state NMR spectrum of a synthetic silicic acid solution (Fig. 5a) and represent the soluble species, disilicic acid (Q^1^) and monosilicic acid (Q^0^). The detection limit of ^29^Si MAS NMR spectroscopy derived from the spectra in Figure 6 b corresponds to a silicic acid (monomer or dimer) concentration of about 1 mM. That means, the presence of significant amounts of silicic acid in whole cells can be excluded. It is concluded that silicic acid is rapidly transformed after uptake, probably to polycondensed species. That means, silicic acid, either in a stabilized or a covalently bound form does not represent large parts of the SSPs in *C. nana* as already observed previously (Gröger et al., 2008).

**Figure 5.**
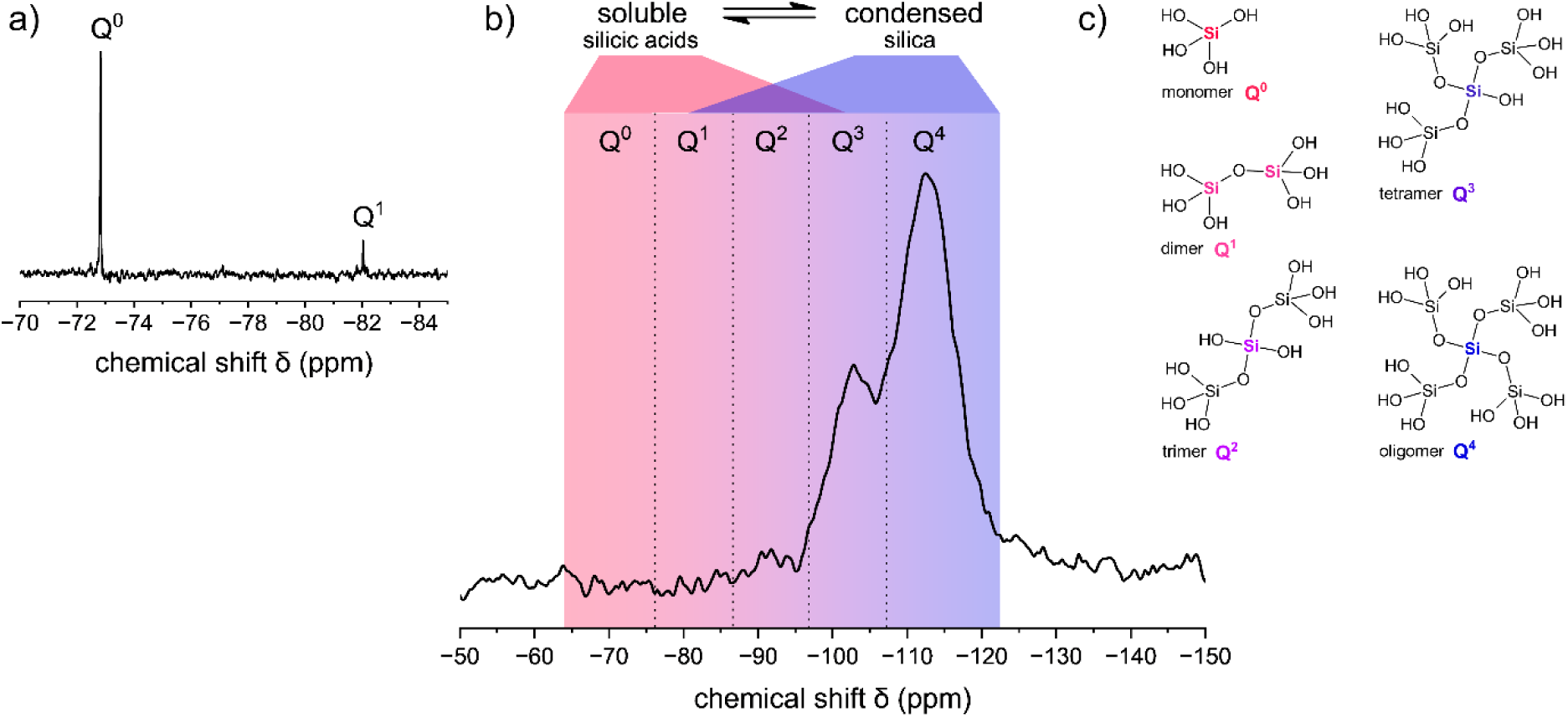
Solid-state ^29^Si MAS NMR spectrum of exponentially grown whole cells (b) with the characteristic Q^n^ group chemical shift regions. For comparison, the liquid-state ^29^Si NMR spectrum of a 4 mM silicic acid solution at pH 9 is inserted (a) showing the signals of the soluble Si species, monosilicic acid (Q^0^) and disilicic acid (Q^1^). These soluble species are not detected in whole cells (region between the dotted lines). c) The intensities of the detected Q^n^ group signals (n = 2,3,4) depend on the degree of silicic acid polycondensation (Belton et al., 2012; McCutchin et al., 2025).

**Figure 6.**
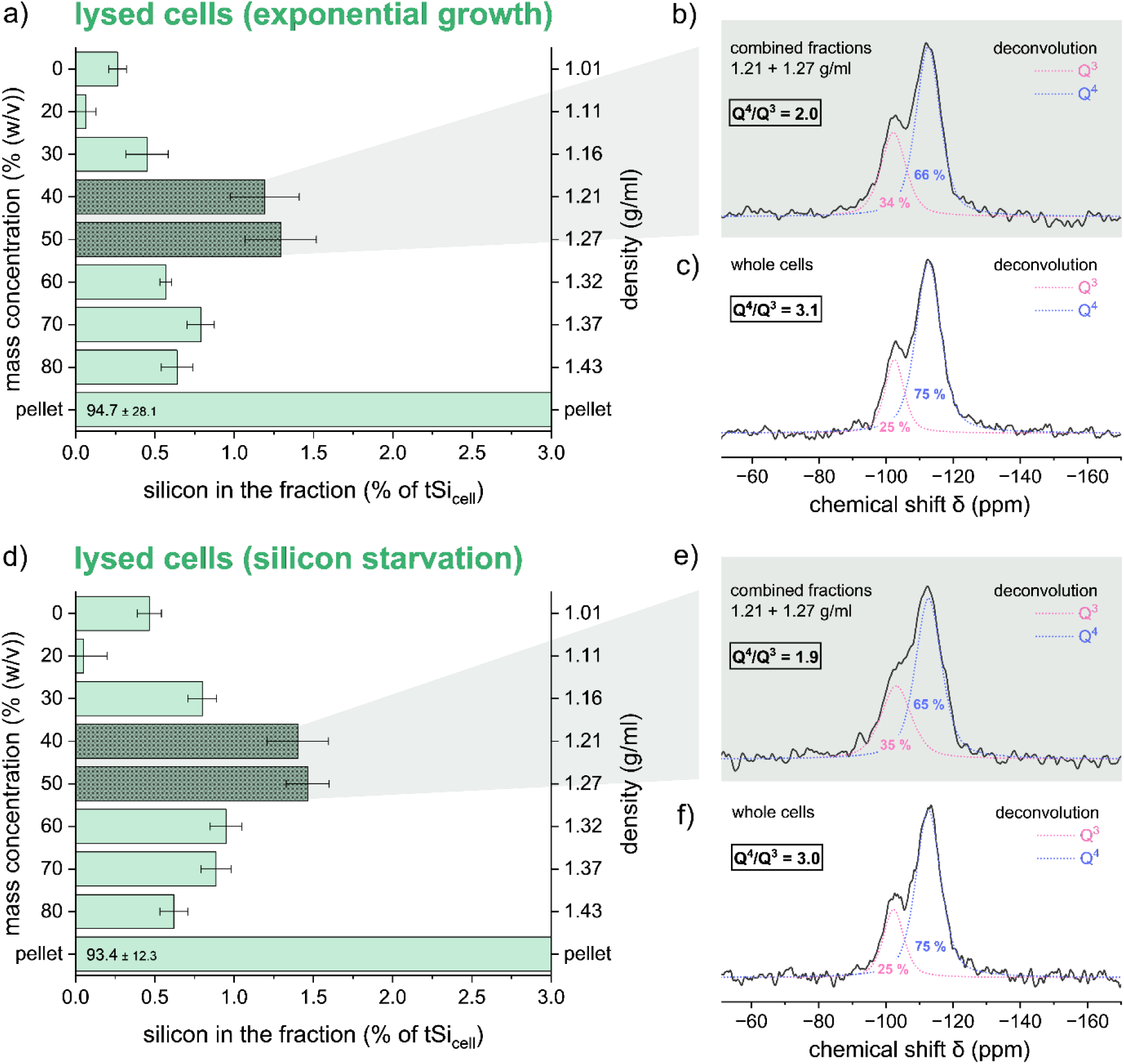
Left: Silicon contents of fractions obtained after density gradient ultra-centrifugation (a,d). Right: Solid-state ^29^Si MAS NMR spectra of whole cells (c,f) and combined fractions (b,e) between 1.21 and 1.27 g/ml (dark green bars). Top: Exponentially grown cells (a,b,c). Bottom: Silicon starved cells (d,e,f). Error bars: 95% CI.

The totally labeled cells were lysed and fractionated by ultracentrifugation as described before (materials and methods). The cytoplasmic silicon content in the density range of 1.01 – 1.27 g/ml was determined to be 2.8 ± 0.6% of tSi_cell_ at exponential growth (Fig.6a) and 3.7 ± 0.3% of tSi_cell_ after silicon starvation (Fig. 6d). Both fully labeled samples came from the same batch and are therefore comparable.

These values are lower compared to the previous separation by ultracentrifugation with unlabeled exponentially growing cells (5.3 ± 0.4% of tSi_cell_). Note that the experimental errors shown in the graphs refer to the silicon quantification method and were derived from the linear regression. These errors do not include the variation of the sample variability and separation protocol. The origin of the large difference is probably the variability between the two different cultivation batches, as diatom cultivation is a complex process and less controlable compared to the separation procedure. This hypothesis is qualitatively supported by the fact that the distribution of silicon over the fractions is similar between the two labeled samples (Fig. 6a and Fig. 6d). Nonetheless, the highest silicon concentrations in the gradient are again observed between 1.21 and 1.27 g/ml as already described above (dark green bars in Fig. 6a,d). These two fractions were combined for NMR measurements of the silicon-containing cytoplasmic constituents (Fig. 6b,e). Spectra of the whole cells were also recorded (Fig. 6c,f) and the integrals of Q^4^ and Q^3^ were obtained by deconvolution using the software *dmfit*. The Q^4^/Q^3^ ratios are provided in the figures. Obviously, the cytoplasmic silica species exhibit a significantly lower degree of silica polycondensation than whole cells which are dominated by cell wall silica. This is in line with results obtained on various species including *C. nana* that already pointed towards pre-condensed silica species in the cytoplasm or the SSPs of diatoms (Bertermann et al., 2003; Gröger et al., 2008).

Comparison of the results for the cultures in exponential growth and after silicon starvation shows no significant differences in the NMR spectra and silicon contents (Fig. 6) except for slightly narrower signals, i.e., better spectral resolution for the sample in exponantial growth and slightly higher cytoplasmic silicon content in silicon starved cells. This implies that the cytoplasmic silicon is not depleted during silicon starvation in agreement with cryoFIB-SEM-EDS studies on *C. nana* (Kumar et al., 2020). On the other hand, the above-described PDMPO labeling experiments clearly show that SSPs are used for biosilica formation during silicon starvation. This should result in a corresponding depletion of the SSPs. It is thus concluded that the clearly observed silica biosynthesis during silicon starvation mainly uses up other SSPs than the species in the cytoplasma separated by our centrifugation experiments. Consequently, there are likely two different silicon pools located in the cell. Based on our observations, we hypothesize the existence of two different silicon pools within the cell. First, the cytoplasmic part of the SSPs. It was detected and characterized through successful separation and following analysis. Cytoplasmic SSPs are characterized by low density (<1.27 g/ml) and are not depleted during silicon starvation (homeostatic). In addition, we postulate the existence of a high-density fraction of SSPs. They cannot be separated using the employed centrifugation techniques because they sediment together with cell wall fragments due to their high density (>1.43 g/mL). That means, we hypothesize the existence of high-density SSPs based on two observations: (i) The significant difference between the results of the PDMPO labeling experiment (SSPs = 34% of tSi_cell_) and the quantification of cytoplasmic silicon (SSPs = 3–10% of tSi_cell_). Comparing these results, the high-density SSPs could potentially account for up to about 30% of tSi_cell_. (ii) Cytoplasmic SSPs are not depleted during silicon-starvation although cell growth continues. It is tempting to speculate that these high-density SSPs represent a distinct cellular (sub)structure. Nevertheless, alternative explanations are also possible. The cells could mobilize silica species which are tightly bound to the frustules or even mobilize cell-wall biosilica for the synthesis of new frustules. Such hypothetical processes could be subject of future studies.

## Discussion

The quantification of the described fluorescence imaging experiments using PDMPO pulse labeling under silicon starvation unequivocally confirms the existence of substantial amounts of silicon stored in SSPs. The use of PDMPO as a proxy for the production of biogenic silica bSiO_2_ provides a relatively simple implementation compared to the expensive and time consuming but very accurate radioisotope labeling with ^32^Si (McNair et al., 2015; Shipe & Brzezinski, 1999). The fluorescence-based method presented here relies on studies of cell cultures grown for a limited time under external silicon starvation. Freshly formed silica must then necessarily be synthesized at the expense of intracellular SSPs. Based on previous work (McNair et al., 2015), the fluorescence intensity of silica-incorporated PDMPO is quantified after pulse labeling. Quantification of this freshly synthesized silica allows to estimate the Si content of the SSPs. To this end, the measured fluorescence intensity from cells under silicon limitation is normalized by the intensity measured for cells in ASW without silicon deficiency. The ratio between the respective fluorescence intensities is a measure for the Si-content in SSPs relative to the total cellular silicon content tSi_cell_ at exponential growth. Species-specific variability of the molar bSiO2: PDMPO incorporation ratio is ruled out by that experimental setup. Five diatom species were examined by this method under laboratory conditions (Tab. S1 and Fig. S1 – S5). For *C. nana*, this method predicts that the SSPs contain up to ca. 34% of tSi_cell_.

It is important to note that this quantification relies on the assumption that the bSiO2 : PDMPO ratio in the formed biosilica does not depend on the external Si concentration in the growth medium. Molar ratios (bSiO₂ : PDMPO) of 2800 ± 700 (Leblanc & Hutchins, 2005) and 2916 ± 708 (McNair et al., 2015) were reported when incorporating PDMPO into biosilica under laboratory conditions. In field studies with a large number of samples that contained whole diatom communities, this relation was confirmed to apply with only slight variations (Leblanc & Hutchins, 2005; McNair et al., 2015). Although, it should be noted that PDMPO incorporation varied by a factor of three among diatom groups (McNair et al., 2015). However, the correlation was only shown to be valid at silicic acid concentrations beyond 3 µM in the growth medium. Below 3 µM, an increasing amount of PDMPO is apparently incorporated into biosilica (McNair et al., 2015) and the incorporation of PDMPO correlates inversely with the silicic acid concentration (Durkin et al., 2013). McNair et al. determined a ratio bSiO_2_ : PDMPO = 912.6 x [Si(OH)_4_] for silicic acid concentrations below 3 µM. This behavior is explained by the change in cell wall composition. Silicon deficiency leads to a higher concentration of biosilica-incorporated organic molecules. It is assumed that PDMPO replaces some of the native organic molecules in the biosilica thus leading to higher concentration of incorporated PDMPO (McNair et al., 2015).

However, the studies of McNair et al. compared the incorporation of PDMPO to the incorporation of ^32^Si, a very sensitive radiomarker (Shipe & Brzezinski, 1999). In short, this method relies on the quantification of biogenic silica based on the measurement of ^32^Si uptake from the growth medium, i.e., bSiO_2_ ∼ ^32^Si incorporation (Krause et al., 2011; McNair et al., 2015). This works very well at high external silicic acid concentrations. But at very low concentrations, pre-existing SSPs in the cells are increasingly mobilized and used for biogenic silica formation in addition to the uptake from the surrounding medium. That means, bSiO_2_ is then not reflected solely by the incorporated ^32^Si but increasingly due to unlabeled silicon originating from mobilized SSPs, i.e., bSiO_2_ ∼ ^32^Si incorporation + unlabeled Si from SSPs. The radioisotope ^32^Si will thus only represent the continuously decreasing, small biosilica fraction formed from external silicic acid taken up under silicon limitation. Given the considerable size of SSPs, mobilization of silicon from SSPs may well explain the apparent increase of the bSiO2 : PDMPO ratio found by the ^32^Si-based method at low external Si concentrations. In contrast, sufficient amounts of silicon are available from the environment at higher external silicic acid concentrations and the SSPs are then not extensively used for biosilica synthesis.

In our experiments using PDMPO labeling instead of ^32^Si, the silicon mobilized from the SSPs will become PDMPO-labeled because the SSP-silicon is transported into the SDV like silicon freshly taken up from the medium and then precipitated whereby PDMPO is incorporated. In summary, the silicon amount of 34% in SSPs relative to tSi_cell_ determined by the fluorescence-based method is definitely an upper bound, but probably even a realistic estimation for the silicon content of SSPs. Given a total cellular Si content of 138 fmol and an average cell volume of 88 fL, this would correspond to a intracellular concentration of ca. 540 mM – far beyond the solubility limit of 2 mM at near-neutral pH (Iler, 1979).

The described separation experiments based on cell lysis by bead beating and subsequent density gradient centrifugation succeeded in separating cytoplasmic silicon. Typical values represent ca. 3 – 10% of tSi_cell_. This corresponds to intracellular concentrations of 50 – 160 mM which also strongly exceeds the solubility limit. However, it is likely that this cytoplasmic fraction only represents part of the SSPs. Furthermore, isopycnic density gradient ultracentrifugation allowed to successfully separate cytoplasmic Si-species of *C. nana* for the first time. The cytoplasmic fraction with a density of 1.01 – 1.27 g/ml shows silicon contents representing about 3 – 5% of tSi_cell_ corresponding to an intracellular concentration of 50 – 80 mM. A maximum is located in the range of 1.21 – 1.27 g/ml. The content of cytoplasmic silicon is stable and appears to be relatively independent of the culture conditions, as shown by a comparison of cells in exponential growth with silicon-starved cells. The spectra of the cytoplasmic silicon fraction are clearly assigned to polycondensed silica species. However, the degree of silica polycondensation is significantly lower than that of the corresponding whole cells which predominantly consist of cell wall biosilica. Consequently, the cytoplasmic silicon consists of silica in pre-condensed form. The presence of such silica was already shown in *C. nana* and other species (Bertermann et al., 2003; Gröger et al., 2008). Here, it could be investigated for the first time in separated form. Note that the presence of significant amounts of siliceous cell walls or their fragments in the cytoplasmic fractions is unlikely for several reasons. First, this is indicated by the sedimentation of the control sample consisting of extracted cell walls. On the other hand, silica fragments produced by cell lysis may sediment differently than the control. However, such biosilica fragments cannot be the dominant part of the detected Si in cytoplasmic fractions because such fragments would exhibit the same degree of silica polycondensation, i.e., the same Q^4^/Q^3^ ratio as the cell walls. This is contradicted by the observations of ^29^Si MAS NMR spectroscopy (see Figure 6) as well as by previous studies on frozen whole cells from synchronized cell cultures (Gröger et al. 2008). This underlines the successful separation of SSPs. Moreover, similar concentrations of intracellular silicon were also reported based on non-invasive techniques like cryoFIB-SEM-EDS (Kumar et al., 2020) which further supports our results.

A remarkable observation is that the silicon content and the NMR spectra of the cytoplasmic silicon remain relatively unchanged when comparing exponentially growing cells with silicon-starved cells. In the first place, this contradicts the assumption that the SSPs are depleted or at least reduced during silicon starvation. On the other hand, the intense growth under silicon starvation observed by the PDMPO labeling experiment clearly demonstrates the extensive use of SSPs and supports their depletion. SSPs could account for up to 1/3 of the tSi_cell_ and thus be significantly larger than previously assumed. This contradiction is resolved by assuming that the synthesis of the cell wall biosilica occurs at the expense of a source other than cytoplasmic silicon. It is concluded that SSPs are consisting of two separate pools: The smaller cytoplasmic low-density fraction (2.8 – 5.3% of tSi_cell_ and <1.27 g/ml) which appears to be stable or homeostatic and a larger high-density fraction (<30% of tSi_cell_ and >1.43 g/ml) which likely represents the actual storage pool.

In order to separate the high-density fraction of the SSPs, the density gradient centrifugation at lower RCF (rate-zonal) is a suitable technique, because separation is mainly based on particle size (Fig. 3). A calculation of the particle migration in the sucrose gradient using the applied parameters (materials and methods) shows that silica particles (2.20 g/ml) are still located in the gradient above the pellet if their radius is below 300 nm (Fig. S6 and App. S1). This is more than a tenth of the average radius of *C. nana* (2440 ± 46 nm). It should be noted that the calculation does not take into account that the particles can deviate from spherical shape, and that the silica particles might have a lower density due to attached organic material or compartimentalization. Both factors would further increase the calculated threshold value for their effective radius. Taking into account the small cell size, it is quite unlikely that such large particles, aggregates or compartments represent the SSPs or a precursor for cell wall formation. Instead, our observations rather point towards the attachment of significant parts of SSPs to the cell walls. It should be noted that a similar hypothesis was already put forward by Lewin in 1954. It was based on the observation that a silicon-starved *N. pelliculosa* culture took up larger amounts of silicic acid under light deprivation. During that period, the cell and valve counts remained unchanged. It was concluded from this observation that no growth took place and silicic acid might be accumulation by attachment, i.e., thickening of the pre-existing siliceous cell walls (Lewin, 1954). Further research should therefore focus on investigating a possible attachment of SSPs and the pre-existing cell walls.

## Conclusion

The knowledge on SSPs in *C. nana* gained by the present study as well as previous work is summarized in Tab. 2. The cytoplasmic silicon content determined here by separation experiments is 3–10% of tSi_cell_ (50-160 mM) and agrees well with the previously determined value of 9% of tSi_cell_ (Reichelt et al., 2023). The resulting cytoplasmic Si concentration of 158 ± 14 mM agrees well with the intracellular silicon concentration of 140 mM determined by cryo-FIB-SEM-EDS experiments (Kumar et al., 2020). This cytoplasmic silicon concentration is, however, significantly smaller than the value of 34% of tSi_cell_ estimated for the SSPs by PDMPO labeling.

**Table 2.** Overview of the methods applied in this investigation and in other studies with the quantitative estimate obtained for the SSPs of *C. nana* in exponential growth. The experimental error is given as 95% CI. * Calculated according to Stokes’ law assuming pure silica particles (Fig. S6-S7 and App. S1). ^†^ Recalculated with the data from the present study (App. S1). ^‡^ Same cultivation batch. ^a^ (Reichelt et al., 2023) ^b^ (Martin-Jézéquel et al., 2000) ^c^ (Thamatrakoln & Hildebrand, 2008) ^d^ (Kumar et al., 2020).

| method | characteristic / information gain | [SSP] / % of tSi<sub>cell</sub> | [SSP] / mM |
|---|---|---|---|
| PDMPO pulse labeling | mobilized during silicon starvation | 34.3 ± 5.7 | 538 ± 96 |
| density gradient centrifugation | particle radius < 300 nm * | 10.1 ± 0.7 | 158 ± 14 |
| isopycnic ultracentrifugation | density < 1.27 g/ml | 5.3 ± 0.4<br>2.8 ± 0.6<sup>‡</sup><br>3.7 ± 0.3<sup>‡</sup> | 83 ± 7<br>44 ± 9<br>58 ± 6 (Si starvation) |
| differential centrifugation<sup>a</sup> | particle radius < 200 nm * | 9 | 141<sup>†</sup> |
| extraction at 95 °C<sup>b, c</sup> | water extractable | 0.5 – 3.5<br>1.3 | -<br>- |
| cryoFIB-SEM-EDS<sup>d</sup> | located in the intracellular space | -<br>- | 140<br>190 (Si starvation) |

Solid-state NMR investigations show that the cytoplasmic silica has a lower polycondensation degree than whole cells, confirming previous reports about the existence of pre-condensed silica species in SSPs. Under silicon starvation, the separated cytoplasmic silicon remains practically unchanged compared to cultures without silicon starvation and the ^29^Si MAS NMR spectra do also not significantly change. Together with the higher amount of SSPs predicted by the fluorescence-based method (34% of tSi_cell_), it seems likely that the cytoplasmic low-density silicon fraction analyzed here only represents a smaller part of the SSPs. The assumption that the presumably much larger fraction of the SSPs consist of large particles, compartments or aggregates seems unlikely in light of particle size calculations based on Stokes’ law. The preferred hypothesis for the location of that missing inseparable high-density part of the SSPs is a connection or association to the cell walls.

## Author contributions

**Eike Brunner**: funding acquisition (lead); project administration (lead); conceptualization (equal); supervision (equal); writing - original draft preparation (equal); writing - review and editing (equal). **Tobias Reichelt**: methodology (lead); formal analysis (lead); visualization (lead); conceptualization (equal); supervision (equal); writing - original draft preparation (equal); writing - review and editing (equal); investigation (equal); validation (equal). **Julia Breyer**: investigation (equal); validation (equal). All authors have read and agreed to the published version of the manuscript.

## Acknowledgments

Financial support by the German Research Foundation (Deutsche Forschungsgemeinschaft, DFG, Project No. Br 1278/35-1) is gratefully acknowledged. The authors gratefully acknowledge Prof. Dr. Thomas Henle (chair of food chemistry – Dresden university of technology) for the possibility to use the ultracentrifuge and Kristina Seliverstova for the instruction and support on that instrument, Dr. Silvia Paasch for supervision and support during NMR experiments, Christiane Leudolph for support in the laboratory work, Yousef Pousti for performing many of the PDMPO labeling experiments and Dr. Susanne Machill for helpful discussions. Figures 1 and 2 were prepared with the software “biorender”.

## Data Availability Statement

The data presented in this study are available on request from the corresponding author.

## Conflicts of interest disclosure

The authors declare no conflict of interest.

## Acknowledgments

Financial support by the German Research Foundation (Deutsche Forschungsgemeinschaft, DFG, Project No. Br 1278/35-1) is gratefully acknowledged.

## Abbreviations

ASW: artificial seawater
ASW-Si: silicon-free artificial seawater
bSiO_2_: biogenic silica
CI: confidence interval
cryo-FIB: cryogenic focused ion beam
EDS: energy-dispersive X-ray spectroscopy
HEPES: 4-(2-hydroxyethyl)-1-piperazineethanesulfonic acid
LOQ: limit of quantification
MAS: magic angle spinning
MBA: molybdenum blue assay
ME: mercaptoethanol
NMR: nuclear magnetic resonance
PDMPO: [2-(4-pyridyl)-5-((4-(2-dimethylaminoethylaminocarbamoyl)methoxy)phenyl)oxazole]
PMSF: phenylmethylsulfonyl fluoride
RCF: relative centrifugal force
SDV: silica deposition vesicle
SEM: scanning electron microscopy
SSPs: silicon storage pools
STVs: silicon transport vesicles
TEM: transmission electron microscopy
tSi: total silicon (amount of substance)
tSi_cell_: total cellular silicon (amount of substance)
[tSi_cell_]: concentration of the tSi_cell_ in the cell volume
tSi_SSP_: total silicon in the silicon storage pools (amount of substance)
[tSi_SSP_]: concentration of the tSi_SSP_ in the cell volume
Q^n^: notation for the different chemical groups of silicon in silicic acids and solid silica
YSZ: yttria-stabilized zirconia

---

**Figure S1.**
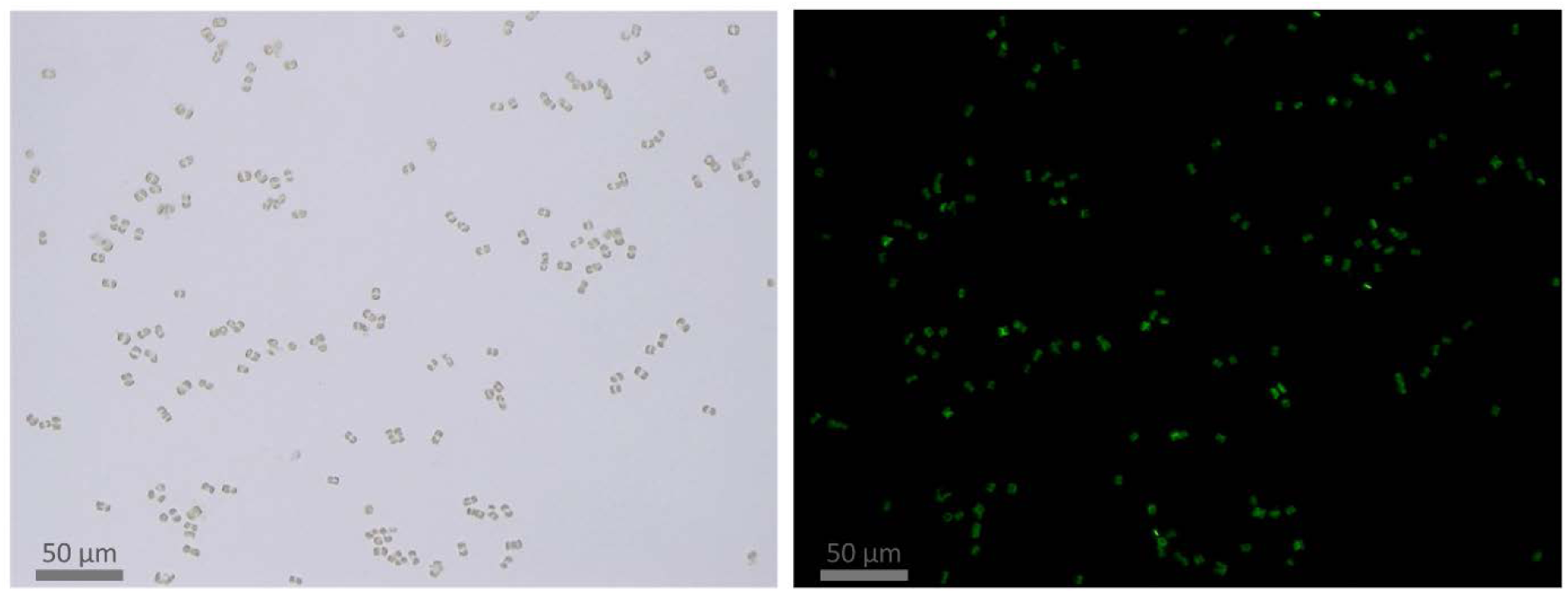
PDMPO pulse labeling of *Cyclotella nana in* silicon-free artificial seawater (ASW-Si). The identic microscopic images are showing a silicon-starved culture of *C. nana* in the bright field channel (left) and the fluorescence channel (right). The fluorescence results from PDMPO incorporation and indicates growth of the cell wall on the expense of the silicon storage pools (SSPs). The cells were pulse labeled with 0.125 µM of PDMPO for 48 h. Note that only one cell of the 151 cells in the image remained unlabeled and has not produced fresh biosilica.

**Figure S2.**
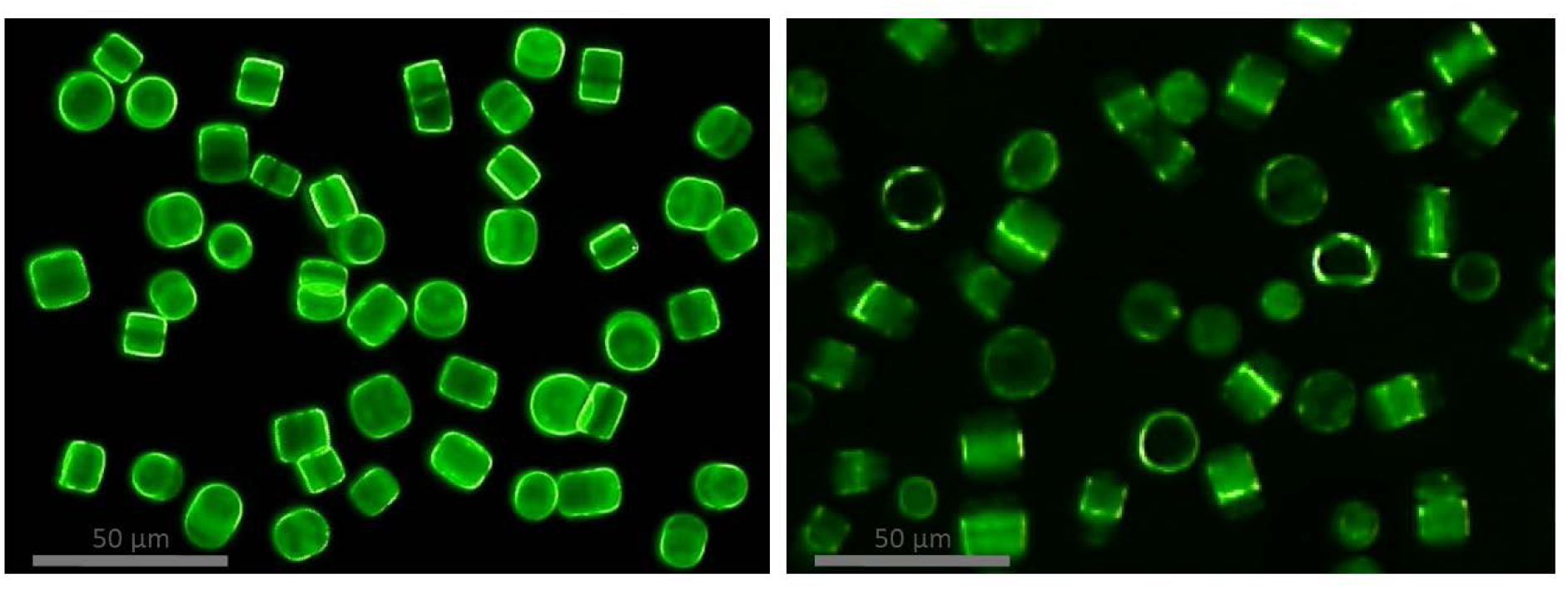
PDMPO labeling of *Conticribra weissflogii*. *C. weissflogii* was totally labeled with PDMPO at exponential growth (left) and pulse-labeled with PDMPO during silicon starvation in ASW-Si (right).

**Figure S3.**
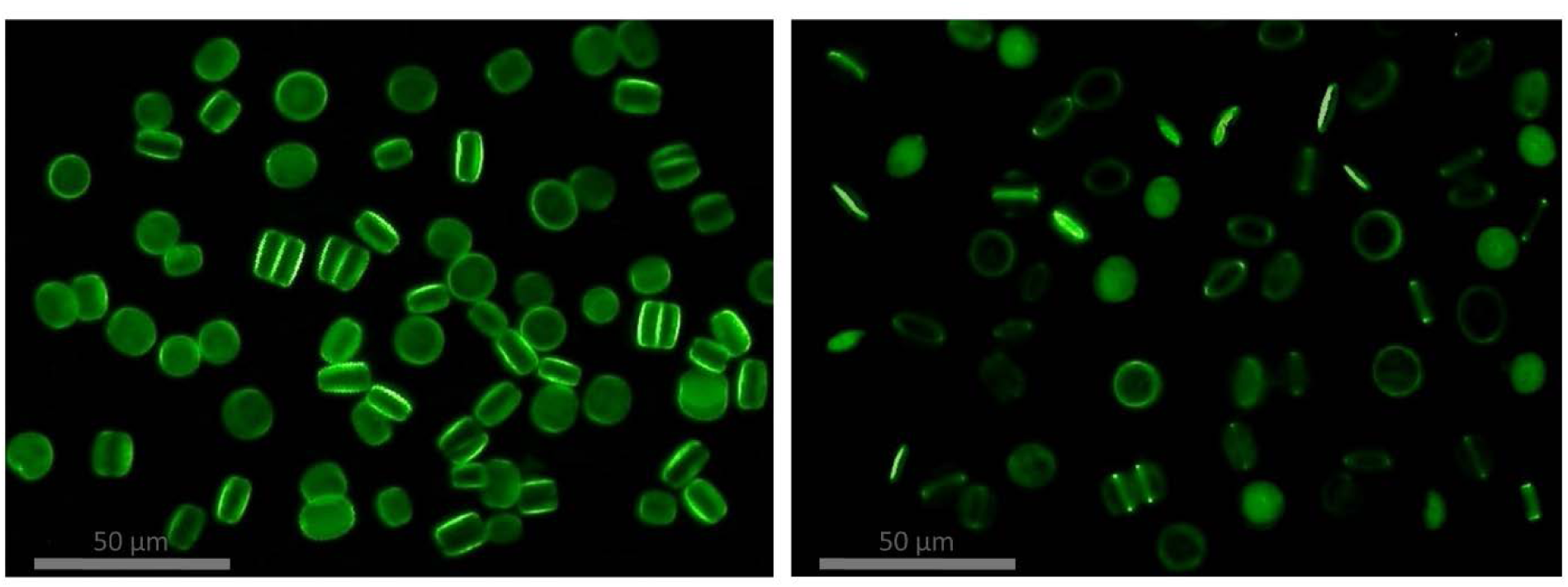
PDMPO labeling of *Cyclotella cryptica*. *C. cryptica* was totally labeled with PDMPO at exponential growth (left) and pulse-labeled with PDMPO during silicon starvation in ASW-Si (right).

**Figure S4.**
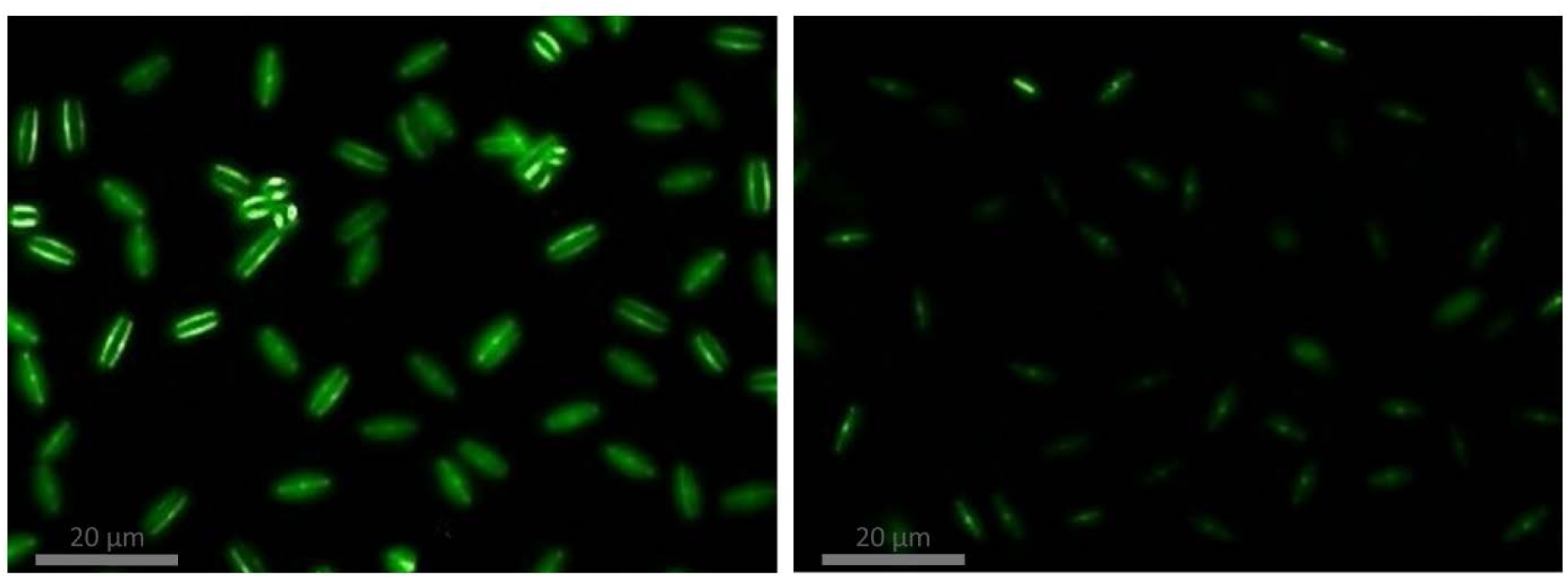
PDMPO labeling of *Navicula pelliculosa*. *N. pelliculosa* was totally labeled with PDMPO at exponential growth (left) and pulse-labeled with PDMPO during silicon starvation in ASW-Si (right).

**Figure S5.**
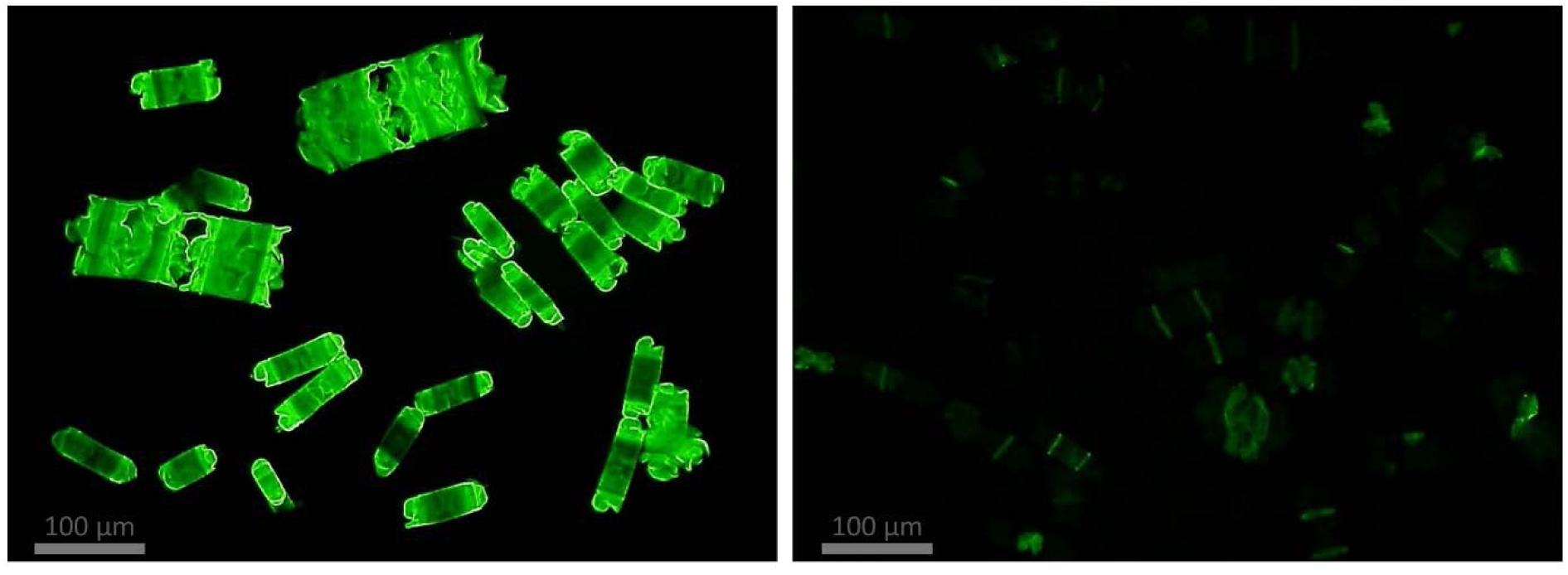
PDMPO labeling of *Biddulphia tridens*. *B. tridens* was totally labeled with PDMPO at exponential growth (left) and pulse-labeled with PDMPO during silicon starvation in ASW-Si (right).

**Figure S6.**
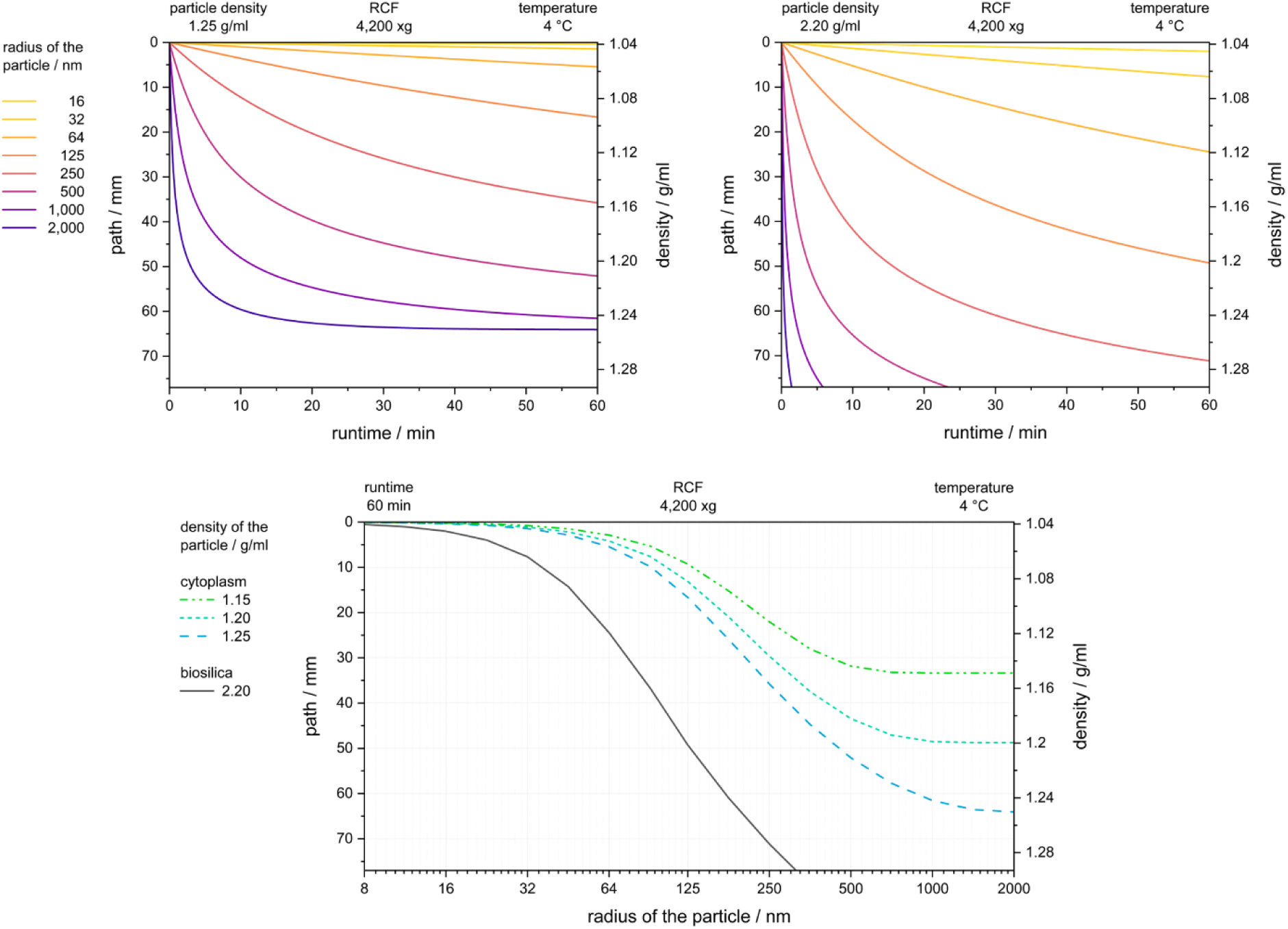
Particle migration in the sucrose gradient. The theoretical particle migration over time is plotted for particles with a density of 1.25 g/ml (top left) and 2.20 g/ml (top right) in the continuous gradient at 4.200 x g. The location of a particle at the end of the 60 min centrifugation run is shown as a function of the radius (bottom). The different particle densities are related to the cytoplasm (color) or biosilica and silica particles (black).

**Figure S7.**
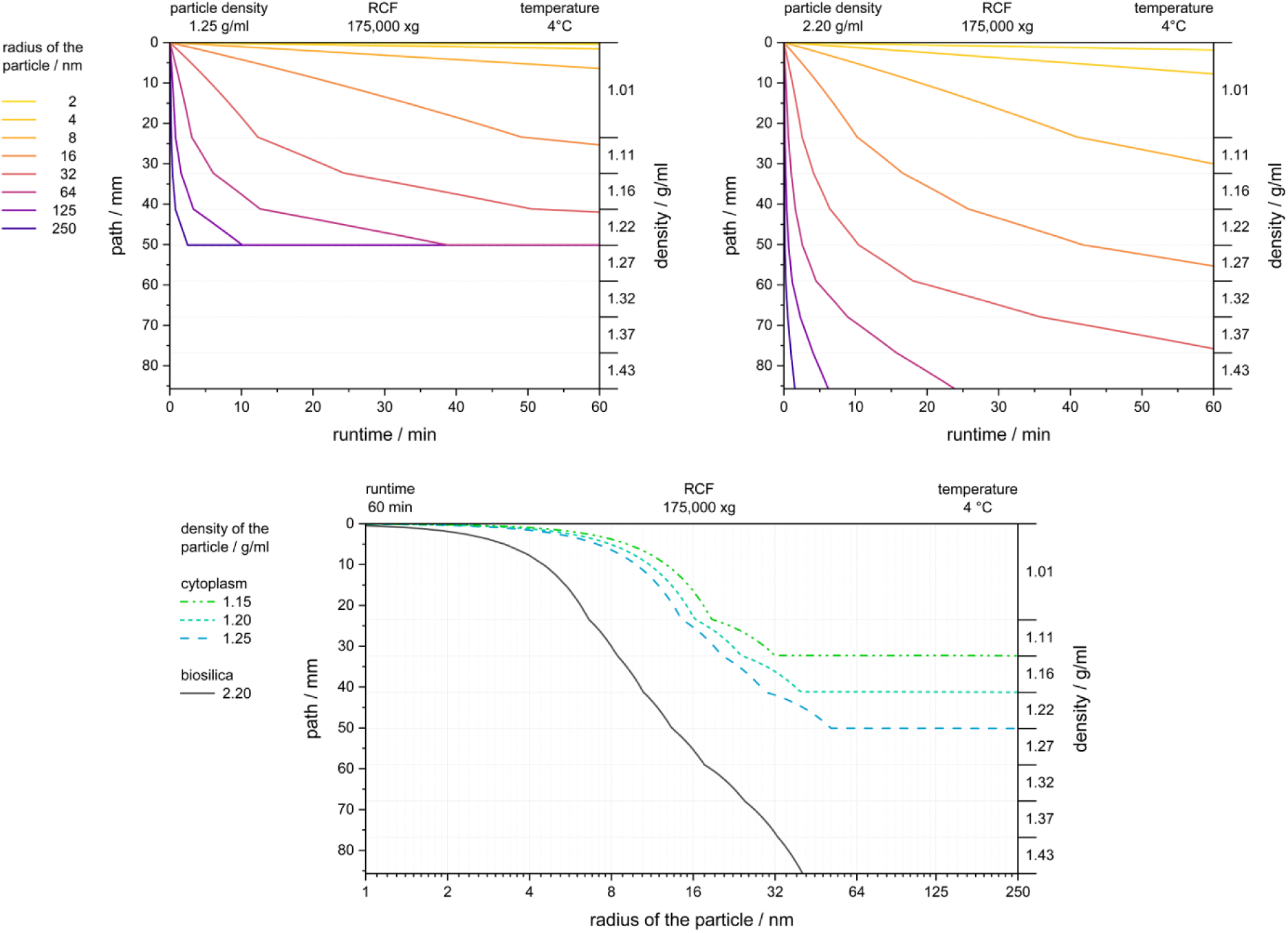
Particle migration in the combined OptiPrep and Nycodenz gradient. The theoretical particle migration over time is plotted for particles with a density of 1.25 g/ml (top left) and 2.20 g/ml (top right) in the stepped gradient at 175,000 x g. The location of a particle at the end of the 60 min centrifugation run is shown as a function of the radius (bottom). The different particle densities are related to the cytoplasm (color) or biosilica and silica particles (black).

**Table S1.** The SSPs of various diatom species were estimated by combined PDMPO pulse labeling and silicon starvation in ASW-Si. The values are given as a fraction of the average total silicon amount of a cell (tSi_cell_). Constant light (48 h) was applied to exclude cell cycle effects. The experiment was reproduced for *C. nana* with results of 35.1 ± 4.8% and 33.5 ± 3.3%. All other experiments were performed once. The experimental error is given as 95% confidence interval.

| species (strain) | SSPs / % of tSi<sub>cell</sub> |
|---|---|
| *Cyclotella nana* (CCMP 1335) | 34.3 ± 5.8 |
| *Conticribra weissflogii* (CCAP 1085/18) | 32.5 ± 4.5 |
| *Cyclotella cryptica* (CCMP 332) | 21.6 ± 2.6 |
| *Navicula pelliculosa* (SAG 1050-3) | 5.5 ± 1.1 |
| *Biddulphia tridens* (unknown) | 4.1 ± 1.8 |

### Appendix S1. Calculation of the particle migration with Stokes’ law

*The movement of a particle is calculated with the law of Stokes. The speed of a particle *ν_p_* is obtained with the following expression.

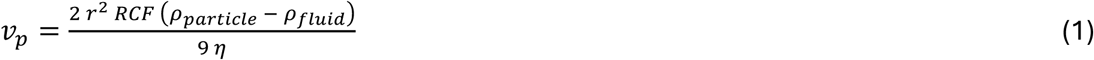

#### Density gradient centrifugation

A continuous gradient of sucrose was used. Therefore, each movement of a particle along the gradient involves a slight change of three parameters in equation (1) which are the density of the medium *ρ_Fluid_*, the viscosity *η* and the rotational centrifugal force *RCF*. Since this correlation is a differential function, the calculation is done with incremental time steps of 1 s. After each step, the three parameters are recalculated with the new position in the gradient. The parameters are expressed as functions of the distance to the rotor centre. The temperature dependent viscosity of the sucrose solution is calculated with the empirical equations presented by Chenlo et al. (Chenlo et al., 2002). The density increase in the gradient is linear due to the use of a gradient mixer. That fact was confirmed by measurements of the refractive index. The upper bound for the particle radius of 300 nm given in Fig. S6 (bottom) is the threshold over which a silica particle with a density of 2.20 g/ml is expected to reach the pellet.

#### Isopycnic density gradient ultracentrifugation

A stepped gradient combining OptiPrep and Nycodenz was used. Therefore, *ρ_Fluid_* and *η* change stepwise in equation (1). The densities were obtained from the data sheets of the manufacturer (Serumwerk, Bernburg, Germany). The viscosities of Nycodenz and OptiPrep at 4°C were estimated using the Andrade equation. Therefore, reported viscosities of OptiPrep and Nycodenz at 20°C and 37°C were used (Eivindvik & Sjogren, 1995).

#### Differential centrifugation

The threshold of the particle radius was calculated with the parameters applied in the cited study (Reichelt et al., 2023), which were a *RCF* of 500 x g, a runtime of 5 min, a path of 20 mm, a density of the medium of 1.01 g/ml and a density of the particles of 2.20 g/ml. The threshold value represents the radius above which a particle is certain to reach the bottom of the tube during centrifugation. Accordingly, particles with radii below this threshold value may remain in the supernatant. It should be noted that at the beginning of the centrifugation the particles are randomly distributed in the solution resulting in random path lengths to the bottom of the tube.

^†^The SSPs of *C. nana* separated by differential centrifugation were estimated to represent 9% of tSi_cell_ (Reichelt et al., 2023). Using the values determined in the present study for tSi_cell_ = 138 fmol and V_cell_ = 88 fL, a recalculated intracellular concentration for the SSPs of 141 mM is obtained. That differs from the value reported in the cited study due to improved techniques.

Figures and tables in the supporting information (entire caption)

## Notes

### Competing Interest Statement

The authors have declared no competing interest.

